# Absolute calibration of ribosome profiling assesses the dynamics of ribosomal flux on transcripts

**DOI:** 10.1101/2023.06.20.545829

**Authors:** Kotaro Tomuro, Mari Mito, Hirotaka Toh, Naohiro Kawamoto, Takahito Miyake, Siu Yu A. Chow, Masao Doi, Yoshiho Ikeuchi, Yuichi Shichino, Shintaro Iwasaki

## Abstract

Ribosome profiling, which is based on deep sequencing of ribosome footprints, has served as a powerful tool for elucidating the regulatory mechanism of protein synthesis. However, the current method has substantial issues: contamination by rRNAs and the lack of appropriate methods to determine overall ribosome numbers in transcripts. Here, we overcame these hurdles through the development of “Ribo-FilterOut”, which is based on the separation of footprints from ribosome subunits by ultrafiltration, and “Ribo-Calibration”, which relies on external spike-ins of stoichiometrically defined mRNA-ribosome complexes. A combination of these approaches measures the absolute number of ribosomes on a transcript, the translation initiation rate, and the overall number of translation events before its decay, all in a genome-wide manner. Moreover, our method revealed the allocation of ribosomes under heat shock stress, during aging, and across cell types. Our strategy transforms ribosome profiling technique from relative to absolute quantification of translation.

## Introduction

Translation is tightly regulated to determine protein abundance in cells. Translational control copes with intra- and extracellular environmental changes. To survey the protein synthesis dynamics in cells, ribosome profiling (or Ribo-Seq) has been developed ^1^ and applied to a wide range of biological contexts ^2–5^. This methodology is based on the protection of mRNA against RNase by ribosomes ^6–8^. Deep sequencing of “ribosome footprints” reveals the positions of ribosomes across the transcriptome at codon resolution, providing a snapshot of the translation status of cells. Since its establishment, ribosome profiling has revolutionized our understanding of protein synthesis. However, there is still an opportunity for technical improvement to overcome the inherent challenges associated with this approach.

A barrier in ribosome profiling experiments has been the extensive contamination of ribosomal RNA (rRNA) fragments that overwhelm the sequencing space. Approaches to limiting this contamination include hybridization to biotin-labeled oligonucleotides followed by immobilization on streptavidin beads ^9–14^ and nuclease-mediated degradation by RNaseH ^15,16^ or duplex-specific nuclease (DSN) ^17^. In addition to direct rRNA fragment targeting, unwanted reads can be reduced through Cas9-mediated degradation after DNA library generation ^12,18–20^. Even with these extensive efforts, however, there is much room to improve rRNA depletion in ribosome profiling.

Another challenge in ribosome profiling is global quantification. While deep sequencing-based analysis can assess relative changes, it is difficult to accurately follow global unidirectional changes in translation in ribosome profiling ^21–23^. To remedy this problem, the supplementation of external standards or “spike-ins” has been employed: the addition of short synthetic oligonucleotides after RNase digestion ^24–27^ or the addition of lysate from an orthogonal species before RNase digestion ^28–31^ has been used for normalization. Similarly, footprints generated by mitochondrial ribosomes could be used for internal spike-ins when translation within organelles is reasonably assumed to be unaffected ^32–36^. However, these methods cannot address the basic question of how many ribosomes reside on each transcript, which hampers the kinetic and stoichiometric understanding of translation in cells.

To overcome these intrinsic difficulties, we developed two key techniques: Ribo-FilterOut and Ribo-Calibration. Ribo-FilterOut uses ultrafiltration to separate ribosome footprints from large and small ribosomal subunits after RNase treatment, deplete fragmented but still complex-assembled rRNAs, and substantially increase the sequencing space for ribosome footprints. Ribo-Calibration employs spike-ins of mol ratio-defined ribosomes associated with mRNA prepared by an *in vitro* translation system; the supplementation of the complex assesses the ribosome numbers on transcripts by data standardization. In combination with other comprehensive techniques, such as ribosome run-off assays and mRNA half-life measurements, Ribo-Calibration revealed the translation initiation speed and the overall number of translation rounds before mRNA decay across the transcriptome. We found that various cellular conditions, including heat shock stress, tissue aging, and cell type differences, may alter the allocation of ribosomes globally and/or in an mRNA-specific manner. By reducing unwanted RNA fragments, our approach provides a versatile platform for absolute quantification of the protein synthesis rate.

## Results

### Separation of ribosome subunits and footprints allows efficient depletion of rRNA in ribosome profiling

One problem in ribosome profiling has been contamination by rRNAs, which overpopulate sequencing libraries and thus reduce the sequencing space for ribosome footprints. For example, in an experiment with HEK293 cells, 92% of the reads were obtained from rRNA [9.2 × 10^5^ reads per million (RPM)], and only 5.4% (0.54 × 10^5^ RPM) of the reads were applicable for downstream analysis (Fig. 1b). Thus, rRNA subtraction by hybridizing oligonucleotides tethered to magnetic beads (such as Ribo-Zero from Illumina) has typically been included in the process of library preparation ^9–14^ (Fig. 1a). This approach reduced rRNA contamination (to 76%) and increased the yield of usable reads (18%) in the library (Fig. 1b). However, a large fraction of rRNAs still escaped subtraction. Thus, an orthogonal approach was needed to improve the output of ribosome profiling.

**Fig. 1:**
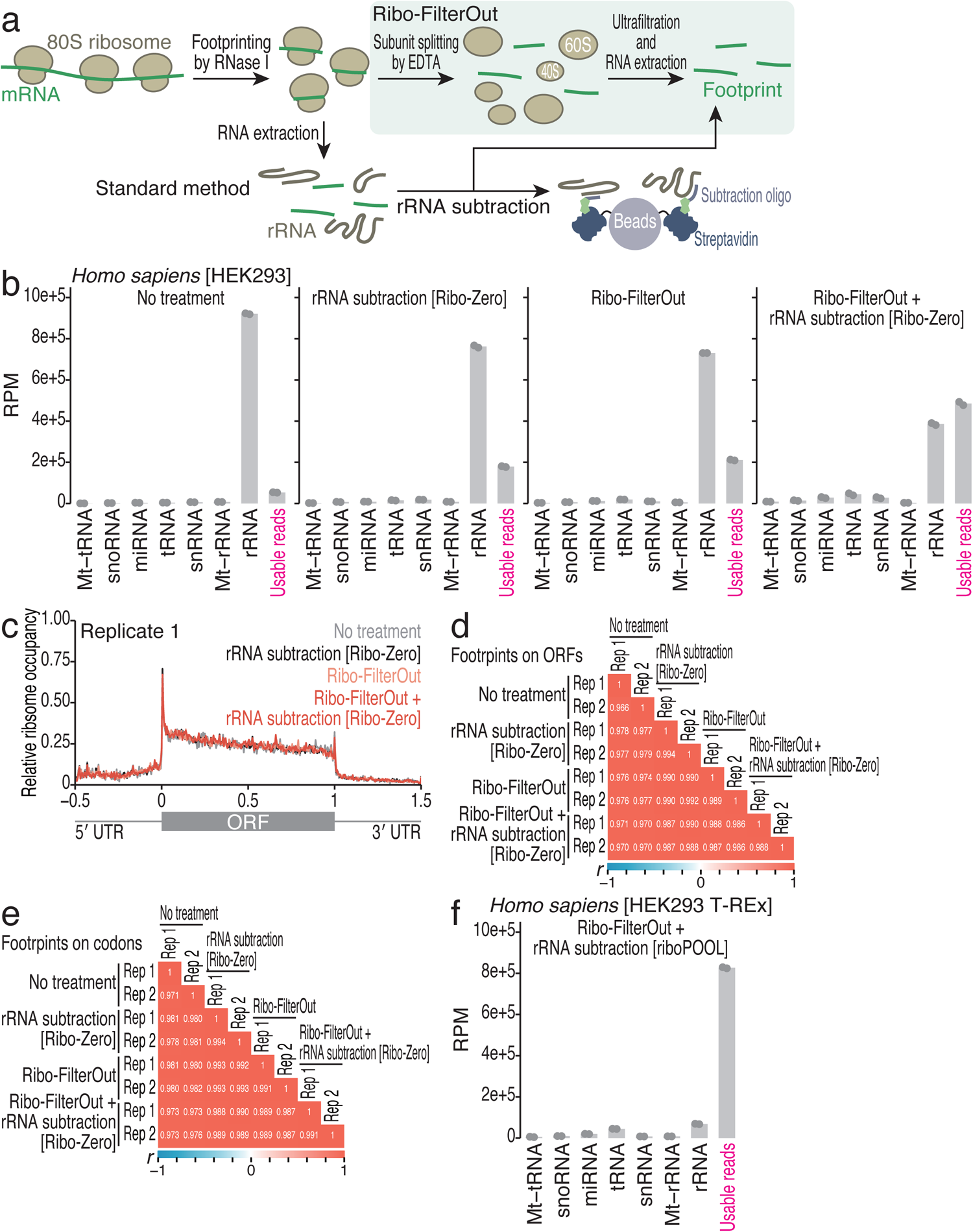
Biochemical separation of ribosome footprints from ribosome subunits reduces contamination by rRNA fragments in ribosome profiling. (a) Schematic of the rRNA depletion methods. The hybridization of biotinylated oligonucleotides and subsequent subtraction on streptavidin beads has been a common strategy (“standard method”). In contrast, Ribo-FilterOut is based on the separation of ribosome footprints from large and small ribosome subunits by ultrafiltration. (b and f) Reads obtained from ribosome profiling were mapped to the indicated RNAs and then to the corresponding genome sequence. The mean (bar) and individual replicates (n = 2, points) are shown. RPM, reads per million mapped reads. (c) Metagene plot of the relative ribosome footprint distribution (ribosome occupancy) along the 5′ UTR, ORF, and 3′ UTR. The length of the ORF was set to 1, whereas the lengths of the 5′ UTR and 3′ UTR were set to 0.5. (d and e) Pearson’s correlation coefficients (*r*) for reads on each ORF (d) or each codon (e). The indicated experimental conditions were compared. The color scales for *r* are shown. Rep, replicate.

Here, we modified the protocol to physically dissociate ribosome footprints from ribosomes (Fig. 1a). In the standard method, ribosomes holding footprints are sedimented by sucrose cushion ultracentrifugation, after which the RNA constituents in the pellet are directly purified. Instead, in the modified protocol, the ribosome pellet is suspended in a buffer containing EDTA, which splits the ribosomes into subunits and releases footprints from the complex (Fig. 1a). The detached footprints are separated from ribosome subunits based on molecular size; the ultrafiltration filter allows small footprints but not macromolecular ribosome subunits containing rRNAs to pass. Indeed, this procedure— termed “Ribo-FilterOut”—eliminated a large amount of RNA fragments, which most likely originated from rRNAs (Supplementary Fig. 1a, b). The remaining ∼30-nucleotide (nt)-long fragments were expected to be ribosome footprints since they disappeared upon EDTA pretreatment, which excludes mRNA from ribosomal protection against RNase digestion (Supplementary Fig. 1a, b).

The application of Ribo-FilterOut improved the performance of ribosome profiling. Ribo-FilterOut increased the number of usable reads (21% in the library) (Fig. 1b). Moreover, the combination of Ribo-FilterOut with rRNA subtraction by hybridizing oligonucleotides (Ribo-Zero) resulted in even greater yields of ribosome footprints (49% in the library) (Fig. 1b), indicating that the application of two different rationales has additive impacts.

Ribosome profiling performed with Ribo-FilterOut can be used to monitor the translatome with limited bias compared with the standard ribosome profiling method. High-quality footprints were obtained in the Ribo-FilterOut-treated samples: the footprint length peaked at 30 and 22 nt (Supplementary Fig. 1c) ^37,38^, 3-nt periodicity was maintained along the open reading frame (ORF) (Supplementary Fig. 1d), and the distribution of reads among the 5′ UTR/ORF/3′ UTR was appropriate (Fig. 1c and Supplementary Fig. 1e, f). Strikingly, the reads mapped to the ORF (Fig. 1d) as well as those assigned to the codons (Fig. 1e) showed high reproducibility between replicates and among the different options for rRNA depletion.

We further tested several conditions to improve the rRNA removal performance in Ribo-FilterOut. We titrated the salt concentration (NaCl) in the resuspension buffer of the ribosome pellet after sucrose cushion centrifugation (Supplementary Fig. 2a). We found that higher salt concentrations (such as 1000 mM) slightly reduced the recovery of usable reads (*i.e.*, increase rRNA fragment contamination). This negative effect of salt could be explained by the drop-off of nicked rRNA fragments from ribosomes. As shown by an earlier study ^39^, RNase treatment may cleave the rRNA within the ribosomes and lead to the drop-off of a subset of ribosome proteins and likely segmented rRNAs. Thus, once rRNA fragments are removed from ribosomes, they pass through the ultrafiltration membrane along with ribosome footprints. On the other hand, rRNA fragments that are retained in ribosomes can be separated from ribosome footprints. Given that high salt may cause the nicked rRNA to dissociate, the use of a moderate concentration of salt (300 mM, throughout this study) during ribosome pellet resuspension may be a better option.

We observed that combining Ribo-Calibration with another rRNA subtraction kit, riboPOOL (siTOOLs Biotech, a system based on probe hybridization and bead subtraction such as Ribo-Zero), improved the performance (Fig. 1f). Ultimately, we were able to use 83% of the reads (∼8.3 × 10^6^ RPM), probably because rRNA probes in the riboPOOL kit may efficiently cover rRNA fragments that escaped from Ribo-FilterOut.

We applied the same strategy to ribosome profiling from other model organisms, such as *D. melanogaster*, *S. cerevisiae*, *A. thaliana*, and *E. coli*. Although the impact of Ribo-FilterOut varied across species, the results in all species, except for yeasts, benefited from the use of Ribo-FilterOut (Supplementary Fig. 2b-e). The differential performance among species should depend on the extent of rRNA digestion by RNase and on the ribosome integrity, as mentioned above. The characteristics of the rRNA sequences, ribosome proteins, and ribosome structures of each species could all affect the performance of Ribo-Calibration.

Overall, the Ribo-FilterOut approach using biochemical separation of footprints from ribosomes was an easily implemented option for high-yield ribosome profiling.

### Ribosome profiling with purified mRNA-ribosome complex spike-ins measures the global shift in protein synthesis

Another challenge in ribosome profiling is to reflect global changes in the protein synthesis rate. To calibrate the ribosome profiling data to measure the net alteration in translation, we used a purified mRNA-ribosome complex with known molarity as a normalization spike-in (termed Ribo-Calibration). The addition of this spike-in to the lysate even before RNase digestion allowed subsequent library preparation procedures without any modification. For this purpose, we harnessed the *in vitro* translation system with rabbit reticulocyte lysate (RRL); mRNAs encoding *Renilla* and firefly luciferases (Rluc and Fluc) were subjected to translation *in vitro,* and then the complexes with two (for Rluc) or three (for Fluc) ribosomes were isolated by sucrose density gradation and ultracentrifugation (*i.e.*, polysome profiling) (Fig. 2a and Supplementary Fig. 3a). We ensured the integrity of the Rluc and Fluc mRNAs in the purified complexes by qPCR of three different amplicons from a transcript (Supplementary Fig. 3b). The addition of the mRNA-ribosome complexes to HEK293 T-REx cell lysate for ribosome profiling enabled us to capture the footprints across the Rluc and Fluc ORFs (Supplementary Fig. 3c), which exhibited high 3-nt periodicity, as was observed for the cellular transcripts (Supplementary Fig. 3d).

**Fig. 2:**
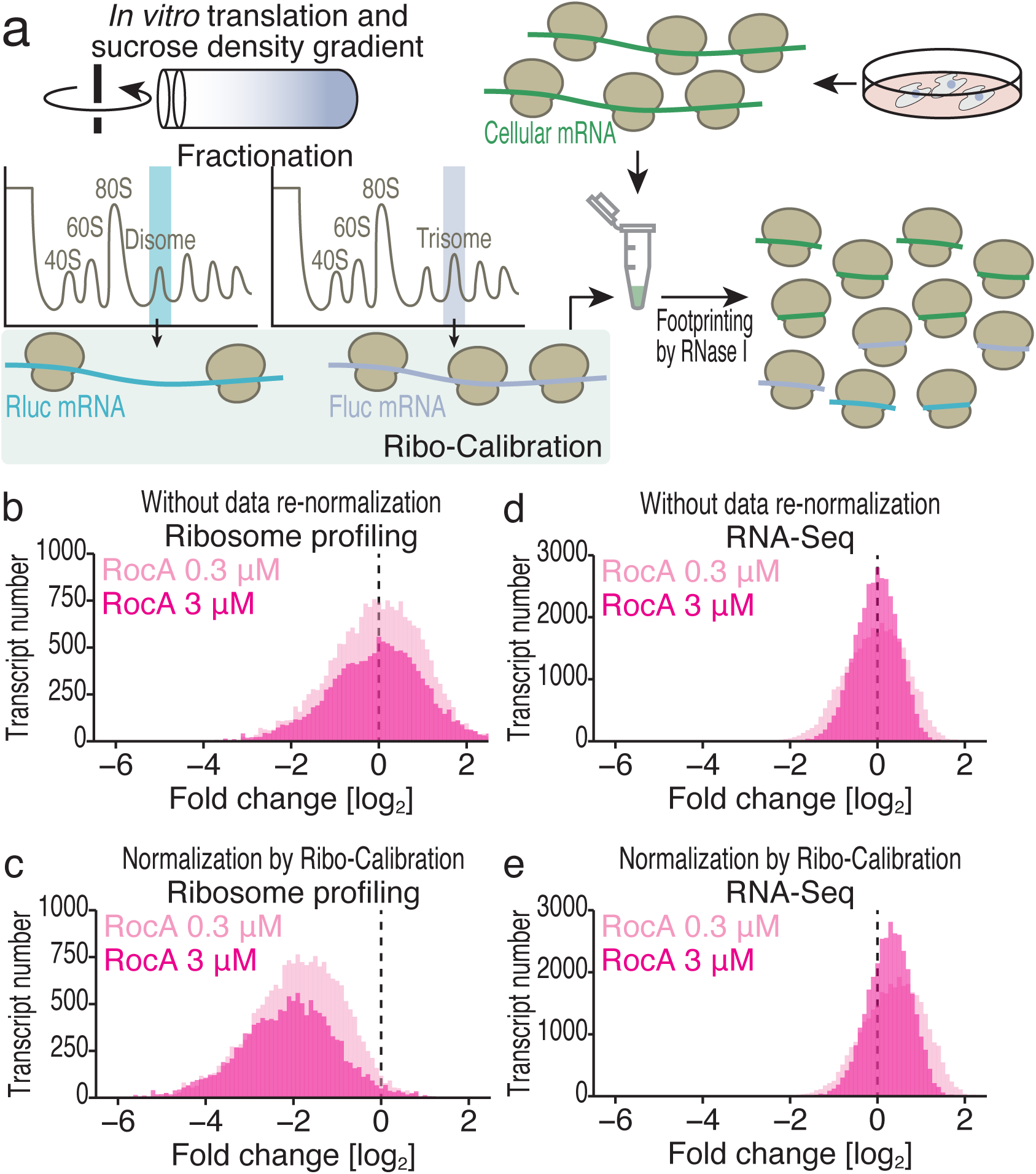
Purified mRNA-ribosome complexes serve as spike-in controls to measure the global change in translation. (a) Schematic of ribosome profiling experiments supplemented with di-ribosome and tri-ribosome-assembled mRNAs as spike-ins. (b and c) Histograms of ribosome footprint fold changes resulting from treatment with 0.3 or 3 μM RocA with (c) or without (b) normalization by Ribo-Calibration spike-ins. (d and e) Histograms of RNA fold changes resulting from treatment with 0.3 or 3 μM RocA with (e) or without (d) normalization by Ribo-Calibration spike-ins.

To test the utility of this spike-in for monitoring global translation, we induced translational repression with rocaglamide A (RocA), a translation inhibitor that targets eukaryotic translation initiation factor (eIF) 4A ^32,34,35,40–42^. For this purpose, we measured the total RNA content in the lysates and used the same amount of material supplemented with an equal volume of purified mRNA-ribosome complexes (Supplementary Fig. 3e).

Renormalization of the data to the reads mapped to the external spike-ins (Supplementary Fig. 3e) allowed us to detect global translation changes induced by RocA (Fig. 2b, c). The same data calibration procedure was applied to simultaneous RNA-Seq of the same materials (Fig. 2d, e). This calculation revealed global accumulation of RNA in response to RocA treatment, implying RNA stabilization due to the loss of translation-coupled RNA decay mechanisms ^43,44^.

In earlier studies ^32,34,35^, ribosome profiling data from RocA-treated cells were normalized to the mitochondrial ribosome (mitoribosome) footprints used as internal spike-ins, assuming that RocA should not cause any translation change in the mitochondria. Although we observed a similar change in global translation with this method and with Ribo-Calibration (Supplementary Fig. 3f), we noted that normalization to the mitoribosome footprint may slightly overestimate the translational repression by RocA (*i.e.*, indicate stronger repression) (Supplementary Fig. 3g). This difference likely originates from the overlooked global change in mRNA abundance as abovementioned. Ribo-Calibration with external spike-ins could prevent this bias.

These data indicated the potential of Ribo-Calibration for data normalization when global protein synthesis is attenuated.

### Ribo-Calibration estimated the absolute number of ribosomes per mRNA across the transcriptome

After the alternative data normalization by Ribo-Calibration, we determined the absolute number of ribosomes per mRNA. Specifically, we assessed the ribosome numbers on each mRNA by determining the stoichiometry between ribosomes and luciferase mRNAs (Fig. 3a and Supplementary Fig. 4a; see Experimental procedures for details). We note that this calculation did not require sample-to-sample comparisons. Rather, data normalization of ribosome profiling and RNA-Seq from the same material and merging those data were key tasks (Supplementary Fig. 4a). The observed ribosome numbers on Rluc and Fluc used for Ribo-Calibration ensured the accuracy of these measurements (Fig. 3b and Supplementary Fig. 4b).

**Fig. 3:**
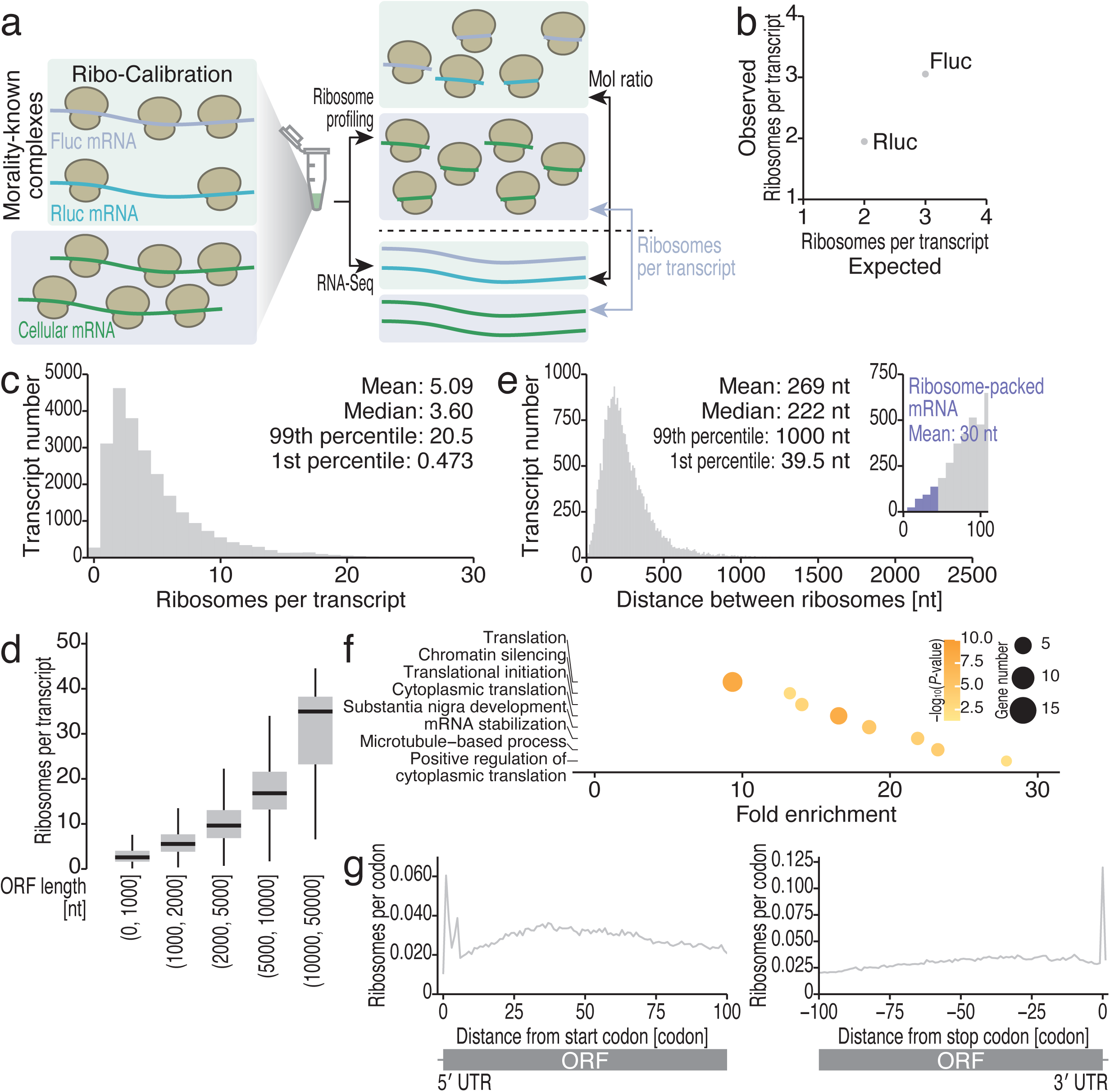
The absolute calibration of ribosome numbers on transcripts. (a) Schematic of the absolute ribosome number calculation by ribosome profiling and RNA-Seq with Ribo-Calibration. (b) Comparison of expected and observed ribosome numbers on Rluc and Fluc transcripts used for Ribo-Calibration. (c) Histogram of the absolute ribosome number on transcripts. (d) Box plots of the absolute number of ribosomes along the ORF length. (e) Histogram of the distance between ribosomes, with the zoomed-in inset highlighting tightly packed transcripts. (f) GO terms enriched in the tightly packed transcripts, defined in e. The color scales for significance and the size scales for gene number in each category are shown. (g) Metagene plots with calibrated ribosome numbers at each position around start codons (left, the first position of the start codon was set to 0) and stop codons (right, the first position of the stop codon was set to 0). In box plots, the median (centerline), upper/lower quartiles (box limits), and 1.5× interquartile range (whiskers) are shown.

We applied this calculation across the transcriptome and observed a high overall correspondence of ribosome numbers on ORFs in replicates (Supplementary Fig. 4c and Supplementary Data 1). Nonetheless, we assessed the errors of the measurements by the Mahalanobis distance in replicates as a benchmark (Supplementary Fig. 4c and Supplementary Data 1); this score was used to identify the outliers quantitatively. We found that mRNAs with low expression (in terms of ribosome profiling reads) generally exhibited high Mahalanobis distance or high variance (Supplementary Fig. 4d). For downstream analysis, we used the mean ribosome numbers on ORFs.

We found a variety of ribosome numbers across ORFs, with 5.09 ribosomes on average (Fig. 3c and Supplementary Data 1). As expected, longer ORFs had more ribosomes (Fig. 3d and Supplementary Data 1). Based on ORF length, the average distance between ribosomes was calculated to be 269 nt (Fig. 3e).

RocA treatment also confirmed the validity of this ribosome molarity calculation. RocA reduced the number of ribosomes globally in a dose-dependent manner (Supplementary Fig. 4e). As expected, RocA greatly increased the distance between ribosomes (Supplementary Fig. 4f). Simultaneously, RocA showed selectivity in repressing a subset of mRNAs ^32,34,35,40–42^ (Supplementary Fig. 4g). The ribosome numbers on high-sensitivity mRNAs were dramatically reduced by RocA, whereas those on low-sensitivity mRNAs were completely resistant (Supplementary Fig. 4h). Thus, absolute ribosome number calibration accurately revealed the ribosome loading rate across transcripts.

### mRNAs with tightly packed ribosomes

We noticed that a subset of mRNAs exhibited an extremely short distance between ribosomes: ∼30 nt, the same length of mRNA occupied by a ribosome (Fig. 3e inset). These “tightly packed” mRNAs were associated with gene categories related to the translation machinery, which requires high expression for active cell growth (Fig. 3f). The ribosome numbers on mRNAs were validated by polysome profiling and subsequent qPCR analysis (Supplementary Fig. 5a-e).

We wondered whether this high density of ribosomes on ORFs can cause ribosome collision. However, a comparison of disome profiling, which captures long footprints generated by two consecutive ribosomes ^19,27,45,46^, revealed that ribosome-packed mRNAs were not significantly associated with ribosome collision (Supplementary Fig. 5f). Thus, although highly ribosome-loaded mRNAs may have a stochastic bump of ribosomes, this may not cause stable ribosome traffic jams that evade the induction of ribosome-associated quality control ^47,48^. The synchronized traversal of ribosomes on the transcript might be the mechanism underlying this phenomenon.

### The absolute number of ribosomes per codon

In addition to mRNA-wise measurements, our method determined the absolute number of ribosomes on each codon. This analysis could assess the probability of ribosome presence in terms of mRNA codons; the averaged ribosome presence along ORFs revealed that start codons may be occupied by ribosomes at ∼6% probability and that stop codons at ∼12.5% probability (Fig. 3g).

Our mRNA-centric analysis of ribosome probability led us to revisit known translation-attenuating codons. *XBP1u* mRNA ^19,49–51^ has been well recognized as a source of ribosome stalling. However, the pause site exhibited only ∼24% occupancy (Supplementary Fig. 5g) due to the relatively low ribosome loading on the transcript (∼1.76 ribosomes per ORF). Recently, upstream ORF 0 (uORF0), which is composed of an ATG start codon and an adjacent stop codon in the 5′ UTR of the *ATF4* mRNA, has been proposed to act as a physical roadblock to prevent ribosome loading downstream ^52^. In our analysis, ∼21% of the uORF0 sequences were covered by ribosomes (Supplementary Fig. 5h), leaving the remaining ∼79% of the uORF0 sequences free of ribosomes. Thus, Ribo-Calibration provided additional information on the probability of ribosome occupancy on mRNAs.

### Measurement of translation initiation rates across the transcriptome

Given that the ribosome number per transcript is a function of the translation initiation rate, translation elongation rate, and ORF length (Supplementary Fig. 6a; see Experimental procedures for details), we reasoned that additional measurement of translation elongation rate may allow us to calculate the absolute translation initiation rates (Supplementary Fig. 6a).

To define ribosome traversal speed, we applied a ribosome run-off assay ^50^; after blocking new ribosome loading onto mRNA by harringtonine (HAR), which inhibits the first round of elongation, preloaded ribosomes were chased and halted by cycloheximide (CHX) over time (Supplementary Fig. 6b). Ribosome profiling under these conditions generated a ribosome-free area downstream of the start codons, extending it over time (Fig. 4a and Supplementary Fig. 6b) ^50^. The regression of the length of the ribosome-free region against time revealed that the elongation rate was 4.08 codon/s on average in HEK293 T-REx cells (Fig. 4b), a similar number to that measured with the same method in mouse ES cells ^50^ and tissues ^53^.

**Fig. 4:**
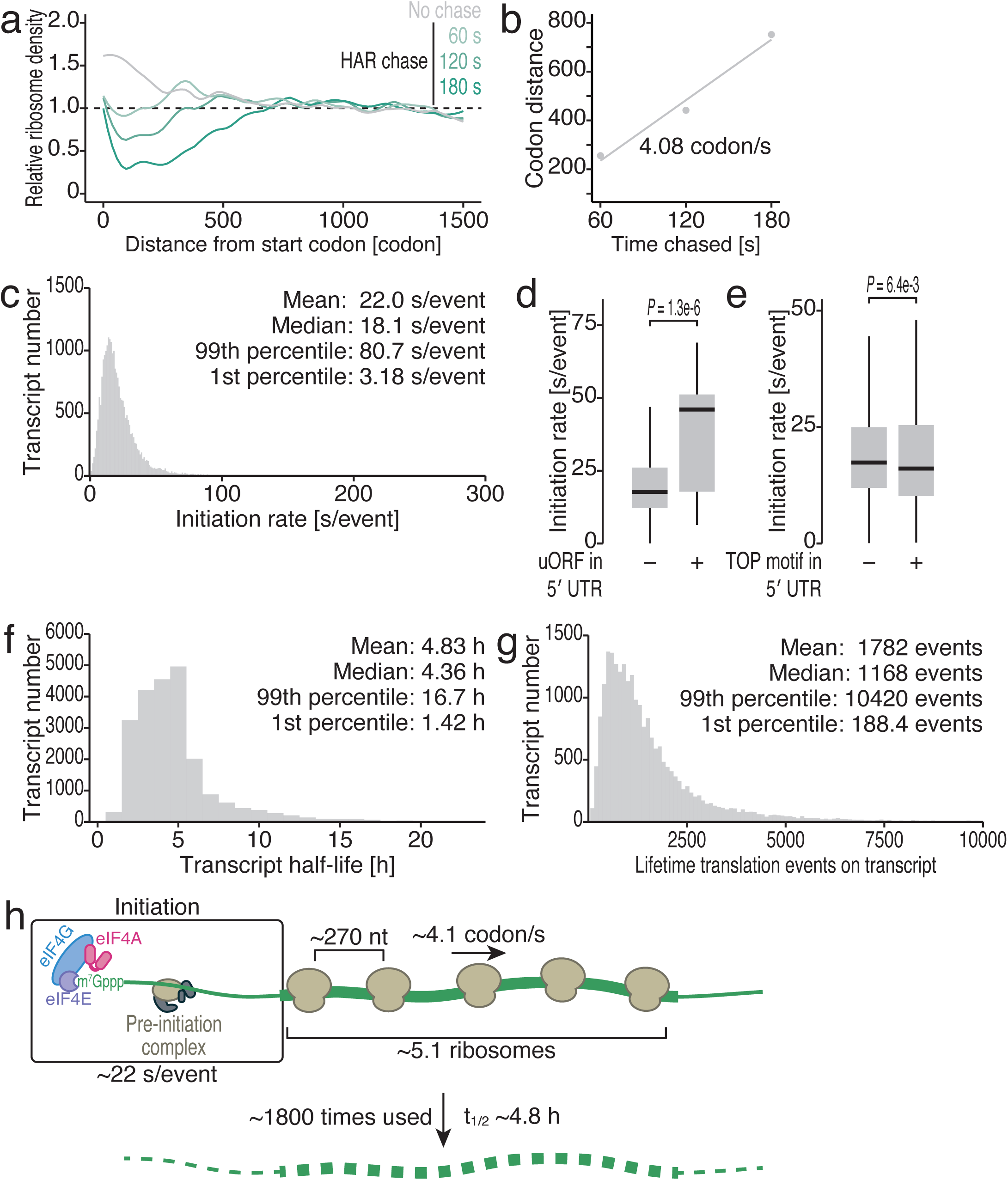
The kinetic parameters determined by Ribo-Calibration. (a) Smoothed metagene plots for a ribosome run-off assay with harringtonine (HAR) chase. (b) The expansion of the ribosome-free area over time by harringtonine chase. Linear regression revealed that the global elongation rate was 4.1 codon/s. (c) Histogram of the translation initiation rate across transcripts. (d and e) Box plots of the translation initiation rate of the mRNAs in the presence or absence of uORFs (defined by ^25^) and TOP motifs (defined by ^98^). The p values were calculated by the Kolmogorov-Smirnov test. (f) Histogram of the transcript half-lives determined by BRIC-Seq. (g) Histogram of the lifetime translation rounds before transcript decay. (h) Schematic summary of the kinetic numbers for translation identified in this study. In box plots, the median (centerline), upper/lower quartiles (box limits), and 1.5× interquartile range (whiskers) are shown.

Then, the translation initiation rate was deduced to be 22.0 s/event on average, but with a wide range across transcripts, from ∼3 s/event to ∼80 s/event (Fig. 4c and Supplementary Data 1). The translation initiation rates of mRNAs are often regulated by *cis* RNA elements. uORFs ^54^ and the 5′ terminal oligo pyrimidine (TOP) motif ^55^ are examples of such negative and positive regulators, respectively. Consistent with this knowledge, we measured slow translation initiation rates for uORF-containing mRNAs (Fig. 4d) and faster rates for TOP motif-bearing mRNAs (Fig. 4e). These observations confirmed our measurements of translation kinetics.

### Lifetime rounds of translation on transcripts

Moreover, given the half-lives of mRNAs, the cycles of mRNA usage for translation could be evaluated. We estimated the mRNA decay rate by 5′-bromo-uridine immunoprecipitation chase–deep sequencing (BRIC-Seq) ^56,57^ (Fig. 4f, Supplementary Fig. 6c, and Supplementary Data 1). Considering the probability of mRNA decay during 5 half-lives (when ∼97% of the mRNA is decayed), mRNAs were subjected to an average of 1782 translation initiations before decay (Fig. 4g and Supplementary Data 1). The lifetime translation events were highly mRNA dependent with a range of ∼55-fold.

Thus, combined with other techniques, our Ribo-Calibration method offered a versatile approach for estimating a variety of kinetic and stoichiometric information related to ribosomal flux (defined as the number of ribosomes initiating translation per unit of time ^58–60)^ across transcriptomes (Fig. 4h).

### The quantification of ribosomal flux in heat shock stress

To evaluate the utility of Ribo-Calibration, we applied this method to translational alterations during physiological stress. For this purpose, we subjected cells to heat shock, a well-recognized stress, to reprogram translation ^61,62^ (Supplementary Fig. 7a). The effects of heat stress and the recovery process were verified via metabolic labeling of newly synthesized proteins using OP-Puro experiments and Western blot analysis of heat shock protein (HSP) 70 family proteins (Supplementary Fig. 7b, c). Ribo-Calibration measured a global decrease in translation caused by heat shock, which peaked at the 1 h time point, and its return to baseline levels during recovery culture (Fig. 5a and Supplementary Fig. 7d). On the other hand, alterations in mRNA abundance were limited (Fig. 5b). Accordingly, the ribosome numbers in the ORFs also followed global translation trends (Fig. 5c and Supplementary Fig. 7e). These data were also consistent with those obtained by the OP-Puro experiments (Supplementary Fig. 7b).

**Fig. 5:**
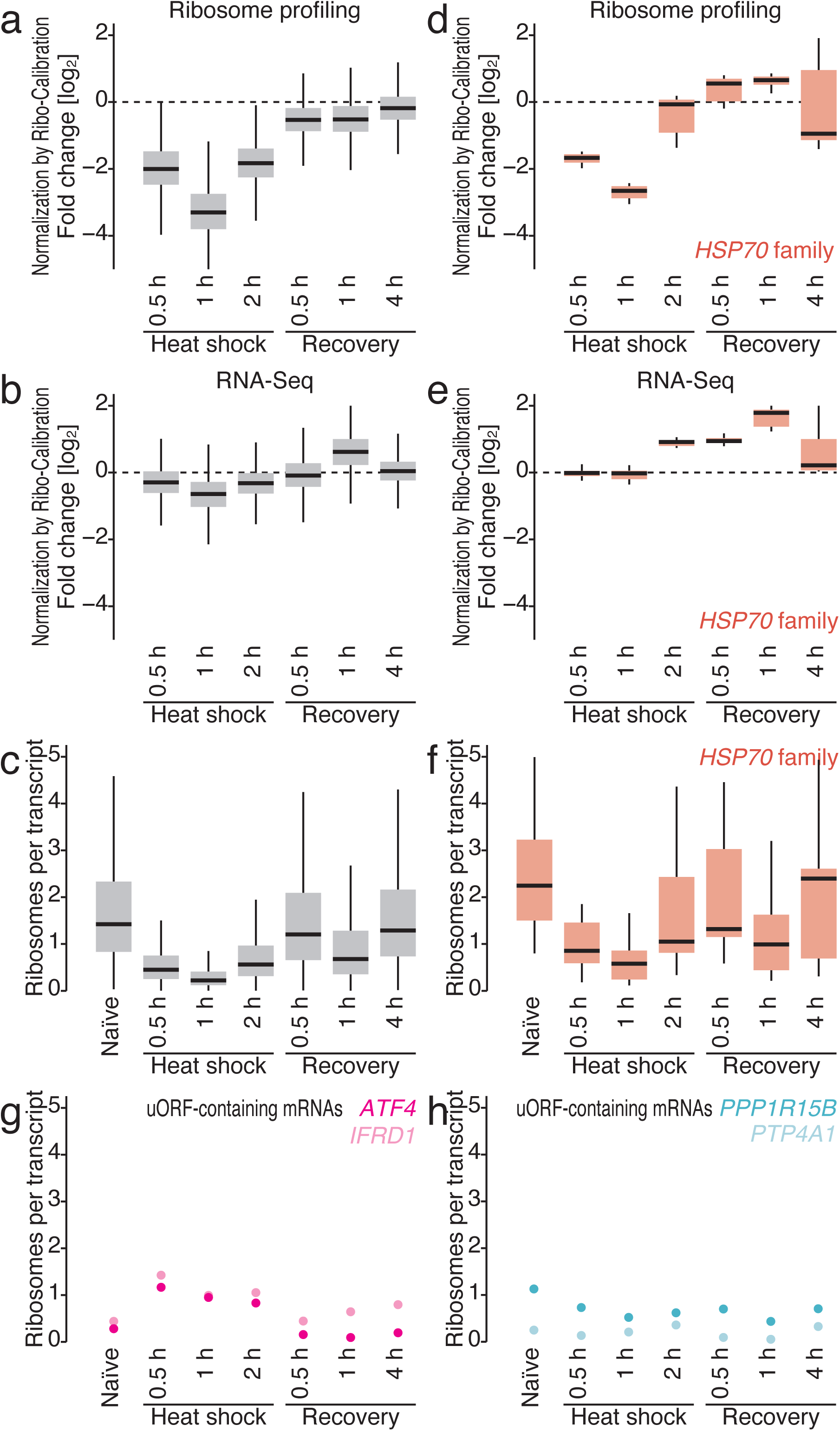
Ribosome flux on transcripts during heat shock stress and recovery. (a and b) Box plots of ribosome footprint fold change (a) and RNA fold change (b) during heat shock and recovery with normalization by Ribo-Calibration spike-ins. (c) Box plots of the absolute ribosome number on transcripts during heat shock and recovery. (d and e) Box plots of ribosome footprint fold change (d) and RNA fold change (e) for transcripts encoding HSP70 family proteins during heat shock and recovery with normalization by Ribo-Calibration spike-ins. (f) Box plots of the absolute ribosome number on transcripts of HSP70 family proteins during heat shock and recovery. (g and h) Dot plots of the absolute ribosome number on transcripts for the indicated genes during heat shock and recovery. In box plots, the median (centerline), upper/lower quartiles (box limits), and 1.5× interquartile range (whiskers) are shown.

Even under the global repression of protein synthesis, heat shock proteins are thought to be preferentially translated to cope with stress ^61,62^. However, the calibrated ribosome profiling data disproved this dogma; during heat shock, the synthesis of HSP70 family proteins was also attenuated (Fig. 5d). Only in the recovery phase, the translation of HSP70 family proteins exceed that in naïve cells, concomitant with transcription induction (Fig. 5e). Ultimately, the number of ribosomes on the ORFs encoding HSP70 family proteins did not surpass that observed in cells before heat shock (Fig. 5f). Thus, the expression of HSP70 family proteins under heat shock stress could be largely explained by transcriptional induction but not translational activation, at least under our conditions.

Another group of mRNAs that are expected to undergo active translation during heat shock consists of uORF-containing mRNAs. Heat shock causes the phosphorylation of eIF2α and an integrated stress response (ISR) ^63^, leading to global translation repression. Simultaneously, the ISR allows the activation of translation from a subset of mRNAs, including those possessing uORFs ^25,63,64^. Among the representative uORF-containing mRNAs ^25^, we observed two types of responses during heat shock: translation was indeed “activated” with increased numbers of ribosomes per ORF (Fig. 5g; exemplified by *ATF4* and *IFRD1*), whereas translation was relatively resistant to global translation reduction; that is, the ribosome numbers on the transcripts remained unchanged (Fig. 5h; exemplified by *PPP1R15B* and *PTP4A1*).

In addition to translation initiation regulation, earlier ribosome profiling studies have indicated ribosome stalling in the early elongation phase (∼65 codons) caused by heat shock ^61^ or by the unavailability of HSP70 family proteins ^65^. Our data recapitulated the higher ribosome occupancy in that region by heat shock when the relative enrichments were considered (Supplementary Fig. 7f). However, the absolute calibration of ribosome numbers across ORF negated such a pileup owing to the global reduction in ribosome loading across the transcriptome (Supplementary Fig. 7g).

Considering these data together, ribosome profiling with Ribo-Calibration provided an accurate view of protein synthesis in stress responses.

### Ribosome flux differences among cell types

We further applied our approach to various cell types, including induced pluripotent stem (iPS) cells and skin cells (melanocytes, fibroblasts, and keratinocytes) (Fig. 6a and Supplementary Fig. 8a-e). We also extended this analysis to tissues from young (3 months) and old (24 months) mouse forebrains (Fig. 6a and Supplementary Fig. 9a-d). We observed differential ribosome numbers on transcripts across different cellular or tissue conditions (Fig. 6a).

**Fig. 6:**
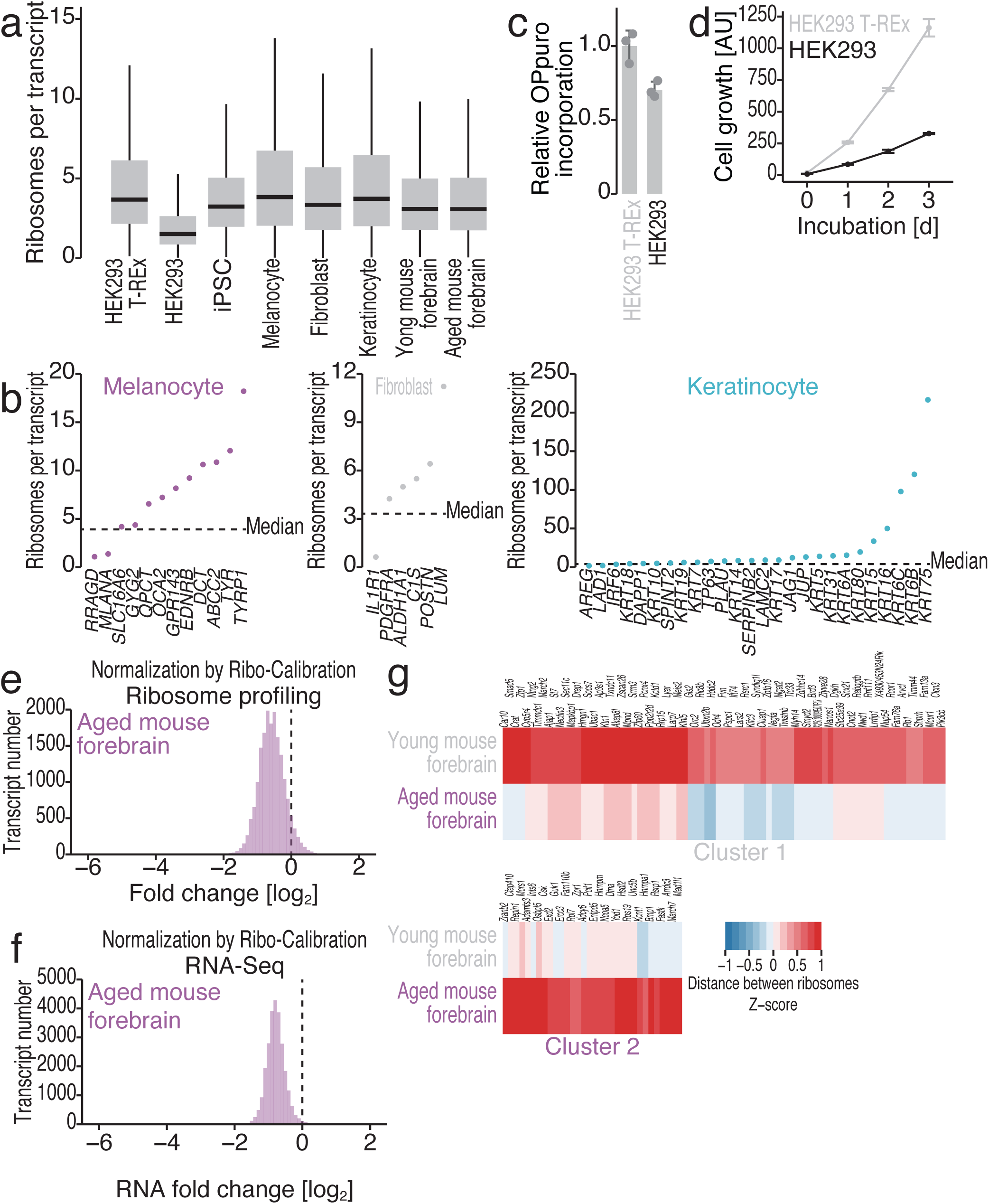
Transcriptome-wide ribosome flux in cell types and aged tissues. (a) Box plots of the absolute ribosome number on transcripts in the indicated materials. (b) Dot plots of the absolute ribosome number on transcripts for the indicated genes in melanocytes, fibroblasts, and keratinocytes. (c) Relative OP-puro incorporation in HEK293 T-REx cells and HEK293 cells. The mean (bar), individual data (n = 3, points), and s.d. (error) are shown. (d) Relative cell growth rate of HEK293 T-REx cells and HEK293 cells. The mean (point) and s.d. (error) from three replicates are shown. (e and f) Histograms of ribosome footprint fold change (e) and RNA fold change (f) in young or aged mouse forebrains with normalization by Ribo-Calibration spike-ins. (g) Heatmap of the distance between ribosomes in young and aged mouse forebrains. The color scales for the Z scores are shown. See Supplementary Fig. 9e for the cluster definition. In box plots, the median (centerline), upper/lower quartiles (box limits), and 1.5× interquartile range (whiskers) are shown.

We noticed that skin cells led to high ribosome fluxes on a subset of mRNAs required for the specialization of the cell ^66,67^: *TYR*, *TYRP1*, and *DCT*, which encode enzymes required for melanin biosynthesis ^68^, in melanocytes; the extracellular matrix protein lumican (LUM), which induces fibrocyte differentiation ^69^, in fibroblasts; and keratin (*KRT75*, *KRT6B*, *KRT6C*, *KRT16*, and *KRT15*) in keratinocytes (Fig. 6b). In addition to genes that define cell functions, differential ribosome loading on a subset of transcripts (here scored by the distance between ribosomes, considering ORF length) was found (Supplementary Fig. 8f-l and Supplementary Data 2).

During our analysis of RocA-mediated translation repression (Fig. 2) and heat shock response (Fig. 5) in HEK293 strains, we noticed that the basal ribosome numbers on the ORFts were very different and that this discrepancy may come from the source/strain of the cells [HEK293 T-REx (from Thermo Fisher Scientific), ∼5.1 ribosomes; HEK293 (from American Type Culture Collection, ATCC), ∼2.3 ribosomes] (Fig. 6a). This difference in ribosome loading reflected the overall protein synthesis, as determined by OP-puro (Fig. 6c), and thus led to differences in growth rate (Fig. 6d).

Therefore, our data indicated that the ribosome flux on individual mRNAs and that at the whole-transcriptome level define cell function and cell proliferation.

### The aged forebrain possesses a lower amount of mRNAs with a constant ribosome loading rate

Consistent with the idea of an aging-dependent reduction in global protein synthesis ^70–72^, we observed declines in overall translation in calibrated ribosome profiling (Fig. 6e). However, this was not reflected in the ribosome numbers on transcripts (Fig. 6a and Supplementary Fig. 9b-d), since the mRNA abundance in the aged forebrain was also decreased (Fig. 6f). Thus, the reduction in global protein synthesis in the aged forebrain was caused by decreased mRNA availability. Even without the overall alteration in ribosome loading, the distance between ribosomes on a subset of transcripts changed during aging (Fig. 6g, Supplementary Fig. S9e-g, and Supplementary Data 3), suggesting that aging induces mRNA-selective dysregulation.

Taken together, these data show that Ribo-Calibration transforms ribosome profiling experiments into a means to measure definite translation with the aid of Ribo-FilterOut for increasing sequencing space.

## Discussion

Here, we showed that the combination of Ribo-FilterOut with the hybridization of oligonucleotides and subsequent rRNA fragment pullout (*i.e.*, Ribo-Zero and riboPOOL) had the best performance. Since a reduction in unwanted reads can be achieved after DNA library generation via Cas9-mediated degradation ^12,18–20^, the application of this strategy may further increase the sequencing depth for usable reads in ribosome profiling. The Ribo-FilerOut approach could be applied to other derivatives of ribosome profiling, such as disome/trisome profiling ^19,27,45,46^ and mitoribosome profiling ^73,74^.

To quantify the global translation change in ribosome profiling data, short DNA or RNA oligonucleotides corresponding to footprint size were used ^24–27^. However, the addition of oligonucleotides was possible only during library preparation, such as after RNase digestion and footprint recovery from ribosomes, and we must assume that no variance in the samples occurred before spike-in supplementation. In contrast, spike-ins of lysates of orthogonal species or purified mRNA-ribosome complexes (*i.e.*, Ribo-Calibration) can be introduced before RNase digestion; thus, this approach is a more suitable experiment design. Moreover, footprints were recovered from the entire spike-in transcripts (Supplementary Fig. 3c), suggesting that this approach provides enough sequence variance of spike-ins and suppresses the bias of normalization that is potentially caused by the limited number of short oligonucleotides ^24–27^. The Ribo-Calibration provides a further advantage for measuring absolute ribosome numbers on transcripts or codons.

Our estimations of the ribosome numbers on transcripts, the distance between ribosomes, and translation initiation rates (22.0 s/event as transcriptome-wide average) aligned very well with those measured by state-of-the-art single-molecule imaging for reporter translation: 10-15 s/event ^75^; 13 and 19 s/event ^76^; 30 s/event ^77^; 29 and 45 s/event ^78^; 17-43 s/event ^79^; 20 s/event ^80^; 54 s/event ^81^; and 13 s/event ^82^. The reported translation initiation speed for one of the most efficient endogenous mRNAs (ovalbumin, 5 s/event) ^83^ also fell within the top 10% (5.7 s/event) (Fig. 4c and Supplementary Data 1). Instead of artificial reporters or the selection of specific mRNAs, our method provides a system-wide assessment of those parameters on the endogenous transcriptome. The independent methods such as a mass spectrometry-based approach that counts the absolute number of newly synthesized proteins will provide complementary validation of these values.

Despite the high correspondence to in-cell kinetics addressed by single-molecule imaging, our genome-wide data may fuel the ongoing debate. Although a recent single-molecule imaging analysis suggested that translation *per se* may trigger mRNA destabilization ^60^, we did not observe a correlation between the translation initiation rate and the mRNA half-life (Supplementary Fig. 6d). The use of average values of mRNAs in cells may make it challenging to determine the fate of individual transcripts, as tracked by single-molecule translation/mRNA imaging.

Another topic of long discussion is a rate-limiting step in protein synthesis: the role of initiation vs. elongation in protein synthesis. Although many studies concluded that the translation initiation dominantly determines the protein synthesis rate in growing cells ^59,75–82,84^. On the other hand, under stressed conditions, the perturbed elongation rate determines the protein production rate ^85–87^. Importantly, our method provides a useful tool to understand kinetic relationships between translation initiation and elongation in wide arrays of cellular situations.

Notably, an earlier study that combined polysome fractionation and RNA-Seq reported a decrease in the number of ribosomes on mRNA with codons longer than 10,000 ^88^. However, this observation may stem from the limitation in the dissection of polysomes larger than 8 ribosomes due to the resolution of the sucrose density gradient experiments. In contrast, our method did not have such a limitation; thus, we found a correlation between ribosome number and length of the ORFs (Fig. 3d).

Considering the cellular contexts that may preferentially allow mono-ribosomes to drive protein synthesis ^89^, the stoichiometry of ribosomes on a transcript may be linked to the function of the cell. The Ribo-Calibration should help to unveil such a connection between the mode of ribosome stoichiometry and phenotype.

### Limitations

Despite the best efforts to avoid the caveats in ribosome profiling techniques, including the unique molecular index (UMI) to suppress biased PCR amplification, the Ribo-Calibration may overestimate the ribosome numbers on ORFs. Even after sucrose cushion to collect the dense complexes including ribosomes, this procedure does not fully exclude the contaminants of footprints from other RNA-protein complexes ^90,91^. Although the fraction may not be huge, careful data interpretation, especially when focusing on individual mRNAs, should be considered.

Another caveat regarding our calculation is the elongation rate that was based on the global average in every codon of the transcriptome. The inclusion of ORF-specific and/or codon-specific elongation rates should be the future direction for the improvement of our technique.

Although this study reported the lifetime translation round, an important parameter, so far, there is currently no alternative method for validating these observations. The development of another strategy to revisit these values is warranted.

## Methods

### Cell lines

HEK293 Flp-In T-REx cells (or HEK293 T-REx; Thermo Fisher Scientific, R78007) and HEK293 cells [American Type Culture Collection (ATCC), CRL-1573] were maintained in DMEM + GlutaMAX-I (Thermo Fisher Scientific) supplemented with 10% fetal bovine serum (FBS, Sigma‒Aldrich) at 37°C and 5% CO_2_.

For RocA treatment, HEK293 T-REx cells were treated with RocA (Sigma– Aldrich) at a concentration of 0.3 μM or 3 μM and incubated for 30 min before cell lysis. Cells treated with 0.1% DMSO (as a solvent for RocA) were used as controls.

For heat shock treatment, HEK293 cells cultured in 6-well plates were transferred to a water bath set at 43°C and incubated for 0.5, 1, or 2 h. For recovery experiments, cells were returned to 37°C and cultured for 0.5, 1, or 4 h.

Human iPS cells (409B2; RIKEN Bioresource Center Cell Bank, HPS0076) were cultured as previously described ^92^.

Human neonatal epidermal melanocytes (HEMn-DP, Thermo Fisher Scientific, C2025C) were maintained in Medium 254 (Thermo Fisher Scientific) containing Human Melanocyte Growth Supplement (Thermo Fisher Scientific). Human neonatal dermal fibroblasts (HDFn, Thermo Fisher Scientific, C0045C) were cultured in Medium 106 (Thermo Fisher Scientific) containing LSGS Supplement (Thermo Fisher Scientific). Human neonatal epidermal keratinocytes (HEKn, Thermo Fisher Scientific, C0015C) were cultured in EpiLife Medium (Thermo Fisher Scientific) with HKGS Supplement (Thermo Fisher Scientific). All primary skin cells were maintained at <80% confluency in a 5% CO_2_/37°C humidified incubator, and the medium was changed every other day. The cells were used between passages 3 and 6.

Fly S2 cells were cultured in Schneider’s *Drosophila* medium (Thermo Fisher Scientific) supplemented with 10% inactivated fetal bovine serum (FBS) (Fujifilm Wako Pure Chemical Corporation) at 27°C.

### Mice

Young and aged C57BL/6J male mice were purchased from CREA Japan Inc. Mice were housed under a regular 12-h light/12-h dark cycle (lights on at 8:00 and off at 20:00) with free access to food and water. All procedures for mouse experiments were conducted in compliance with the Ethical Regulations of Kyoto University and performed under protocols approved by the Animal Care and Experimentation Committee of Kyoto University.

### Yeast

Yeast cells (BY4741) were cultured in a nutrient-rich liquid medium supplemented with Difco YPD Broth (BD Biosciences) at 30°C.

### Plants

Wild-type *Arabidopsis thaliana* (Col-0) seeds were sterilized using sodium hypochlorite and washed 5 times in sterile water. The seeds were sown on Murashige and Skoog’s half-strength solid medium (1/2 MS medium) supplemented with 3% sucrose and incubated in the dark at 4°C. After 2 d, the plants were grown at 23°C under a 12-h light/12-h dark cycle for 13 d. The seedlings were collected under light conditions, frozen in liquid nitrogen, and stored at −80°C.

### Bacteria

*E. coli* MG1655 cells were grown in Miller LB medium (nacalai tesque) at 37°C.

### Preparation of spike-in Rluc-disome and Fluc-trisome

PCR amplification was performed using the psiCHECK2-EIF2S3 ^32^ plasmid as a template with the following forward and reverse primers: Rluc, 5′-TAATACGACTCACTATAGG-3′ and 5′-CACACAAAAAACCAACACACAG-3′; Fluc, 5′-ACTTAATACGACTCACTATAGGAAGCTTGGCATTCCGG-3′ and 5′-TGTATCTTATCATGTCTGCTCGAAG-3′. These DNA fragments were subjected to *in vitro* transcription using a T7-Scribe Standard RNA IVT Kit (CELLSCRIPT). The RNAs were then capped and polyadenylated using a ScriptCap m^7^G capping system (CELLSCRIPT), a ScriptCap 2′-*O*-Methyltransferase Kit (CELLSCRIPT), and an A-Plus poly(A) polymerase tailing kit (CELLSCRIPT).

Rluc mRNA and Fluc mRNA were incubated with rabbit reticulocyte lysate (RRL, Promega) to generate the mRNA-ribosome complex. The reaction mixture consisted of 50 μl of lysate, 30 μl of H_2_O, 10 μl of 500 nM mRNA, and 10 μl of premix [100 μM amino acid mixture without methionine (Promega), 100 μM amino acid mixture without leucine (Promega), and 1 U/ml ScriptGuard RNase inhibitor (CELLSCRIPT)]. The mixture was incubated at 30°C for 0.5 h (Rluc mRNA) or 1 h (Fluc mRNA) and mixed with an equal volume of 2× lysis buffer (40 mM Tris-HCl pH 7.5, 300 mM NaCl, 10 mM MgCl_2_, 2% Triton X-100, 2 mM DTT, 200 µg/ml cycloheximide, and 200 µg/ml chloramphenicol).

The lysates were loaded directly onto a 10-50% sucrose gradient in SDG buffer (20 mM Tris-HCl pH 7.5, 150 mM NaCl, 5 mM MgCl_2_, 1 mM DTT, 100 µg/ml cycloheximide) and ultracentrifuged at 35,300 rpm and 4°C for 2.5 h using a Himac CP80WX ultracentrifuge (Hitachi) with a P40ST rotor (Hitachi). The gradients were fractionated with continuous measurement of absorbance at 260 nm using a Triax flow cell (BioComp) and a micro collector (ATTO). Fractions containing disomes on Rluc mRNAs or trisomes on Fluc mRNAs were pooled, resulting in a total volume of 1.1 ml for each. These combined fractions were quickly frozen using liquid nitrogen and stored at −80°C.

The integrity of the Rluc and Fluc mRNAs in the isolated complex was validated via qPCR with 3 different amplicons. Total RNA from the lysate supplemented with the purified mRNA-ribosome complexes was extracted using TRIzol LS reagent and a Direct-zol RNA MicroPrep Kit and treated with a TURBO DNase (Thermo Fisher Scientific). cDNAs were synthesized using the ReverTraAce qPCR RT Kit (TOYOBO), following the manufacturer’s protocols. RT‒qPCR analyses were conducted using an iTaq Universal SYBR Green Supermix Kit (Bio-Rad) and a CFX Connect Real Time System (Bio-Rad). The titrated amount of bare mRNAs used for *in vitro* translation was used for data calibration. The following primers were used: Rluc amplicon 1, 5′-ATGGCTTCCAAGGTGTACGA-3′ and 5′-TCCGATCAGATCAGGGATGA-3′; Rluc amplicon 2, 5′-TCGTCCATGCTGAGAGTGTC-3′ and 5′-CTAACCTCGCCCTTCTCCTT-3′; Rluc amplicon 3, 5′-CAACTACAACGCCTACCTTC-3′ and 5′-TTACTGCTCGTTCTTCAGCA-3′; Fluc amplicon 1, 5′-ATGGCCGATGCTAAGAACAT-3′ and 5′-TGTAAATGTCGTTAGCAGGG-3′; Fluc amplicon 2, 5′-TTTCGACAGGGACAAAACCA-3′ and 5′-GCAGGGCAGACTGAATTTTG-3′; and Fluc amplicon 3, 5′-CAAGTACAAGGGCTACCAGG-3′ and 5′-CCGCCTTTCTTAGCCTTGAT-3′.

### Ribosome profiling and RNA-Seq

Ribosome profiling libraries were prepared as previously described ^12^ with some modifications. The libraries prepared in this study are summarized in Supplementary Data 4.

#### Lysate preparation

HEK293 T-REx cells were cultured in a 10-cm dish in preparation for RocA treatment, and HEK293 cells were cultured in 6-well plates for heat shock treatment. The cells were then lysed in 500 µl and 100 µl of lysis buffer (20 mM Tris-HCl pH 7.5, 150 mM NaCl, 5 mM MgCl_2_, 1% Triton X-100, 1 mM DTT, 100 µg/ml cycloheximide, and 100 µg/ml chloramphenicol) and treated with 25 U/ml TURBO DNase (Thermo Fisher Scientific) ^12^.

Human skin cells cultured in 6-well plates and iPS cells cultured in 12-well plates were lysed in 100 μl of lysis buffer and treated with 25 U/ml TURBO DNase ^12^.

The mouse forebrain was dissected on ice and disrupted using a loose-fitting Dounce homogenizer in 3 ml of ice-cold homogenizing buffer (320 mM sucrose and 5 mM HEPES-KOH pH 7.4). The homogenate was treated with an equal volume of 2× lysis buffer and incubated for 10 min on ice with 25 U/ml TURBO DNase.

S2 cells were seeded in a 10-cm dish, cultured for 2 d, treated with 100 μg/ml cycloheximide and 100 µg/ml chloramphenicol, and then collected by immediate centrifugation at 1,000 × *g* for 3 min. The cell pellet was washed with ice-cold PBS containing 100 μg/ml cycloheximide and 100 µg/ml chloramphenicol, and then lysed with 400 μl of lysis buffer and treated with 25 U/ml TURBO DNase.

For yeast, cells were collected by filtration, snap-frozen in liquid nitrogen with ice grains of lysis buffer, and pulverized using a Multi-Beads Shocker with a precooled chamber and ball with liquid nitrogen ^93^. The cell lysate was then centrifuged at 3,000 × *g* for 5 min to remove cellular debris. The supernatant was transferred to a new tube and treated with 25 U/ml TURBO DNase.

Snap-frozen *Arabidopsis thaliana* seedlings and ice grains of 400 μl of plant lysis buffer (100 mM Tris-HCl pH 7.5, 40 mM KCl, 20 mM MgCl_2_, 1 mM DTT, 300 μg/ml chloramphenicol, 100 μg/ml cycloheximide, and 1% Triton X-100) were pulverized using a Multi-Beads Shocker with a precooled chamber and ball with liquid nitrogen ^94^. The lysate was clarified by centrifugation at 3,000 × *g* for 5 min at 4°C and treated with 25 U/ml TURBO DNase for 10 min on ice.

After DNase treatment, the eukaryotic cell lysate was clarified by centrifugation at 20,000 × *g* for 10 min at 4°C.

*E. coli* cells grown overnight at 37°C in LB medium were diluted 1:100 in fresh LB medium and grown at 37°C until the optical density at 600 nm (OD_600_) reached 0.4-0.5 ^95^. Ten milliliters of culture were then collected by direct freezing in liquid nitrogen. The frozen samples and 5× bacterial lysis buffer (100 mM Tris-HCl pH 7.5, 750 mM NH_4_Cl, 50 mM MgCl_2_, 25 mM CaCl_2_, 5 mM DTT, 5% Triton X-100, 10 mg/ml capreomycin, and 500 µg/ml tigecycline) were cryo-milled using a Multi-Beads Shocker [Yasui Kikai, MB2200(S)] with a precooled chamber and ball with liquid nitrogen. After thawing at room temperature, the supernatant of the crude lysate was obtained by centrifugation at 9,000 *× g* and 4°C for 10 min. To enrich ribosome fractions, the supernatant was then subjected to ultracentrifugation with sucrose cushion buffer (20 mM Tris-HCl pH 7.5, 150 mM NH_4_Cl, 10 mM MgCl_2_, 1 mM DTT, 1% Triton X-100, 2 mg/ml capreomycin, 100 µg/ml tigecycline, 20 U/ml SUPERase•In RNase Inhibitor, and 1 M sucrose) at 100,000 rpm (543,000 *× g*) and 4°C for 1 h with a TLA110 rotor (Beckman Coulter) and an Optima MAX-TL Ultracentrifuge (Beckman Coulter). Ribosome pellets were resuspended in 1× bacterial lysis buffer.

Total RNA concentrations in the lysates were measured with a Qubit RNA Broad Range Assay Kit (Thermo Fisher Scientific). The lysates were flash-frozen with liquid nitrogen and stored at −80°C.

#### Ribo-Calibration

The lysate of HEK293 T-REx cells containing 12 µg of total RNA was adjusted to a volume of 338 µl using lysis buffer. Then, 6 µl of Rluc-disome spike-in and 6 µl of Fluc-trisome spike-in were added to the lysate to make a 350 µl lysate solution. For ribosome profiling, 290 µl of the lysate solution (corresponding to 10 µg of total RNA) was incubated with 10 µl of 10 U/µl RNase I (Lucigen) for 45 minutes at a temperature of 25°C. For RNA-Seq, the remaining 50 µl of the lysate solution was directly subjected to RNA purification with TRIzol LS reagent (Thermo Fisher Scientific) and a Direct-zol RNA MicroPrep Kit (Zymo Research).

For heat shock experiments in which HEK293 cells were used, the lysate containing 7 µg of total RNA was adjusted to 350 µl.

For human skin cells and iPS cells, the volume of the lysate was adjusted to 350 µl from an initial 100 µl by lysis buffer.

#### Ribo-FilterOut and rRNA subtraction

After RNase digestion and sucrose cushion ultracentrifugation (at 100,000 rpm for 1 h at 4°C with a TLA110 rotor and an Optima MAX-TL Ultracentrifuge), the ribosome pellet was resuspended in 150 μl of resuspension buffer (20 mM Tris-HCl pH 7.5, 300 mM NaCl, 5 mM EDTA, 1% Triton X-100, 1 mM DTT, and 20 U/ml SUPERase•In). For the salt titration, 300 mM NaCl was replaced with 150 mM or 1000 mM NaCl. The mixture was then transferred to the filter cup of an Amicon Ultra 0.5 ml Ultracel 100K centrifugal filter (Millipore) and centrifuged at 14,000 × *g* at 4°C for 10 min. The flowthrough in the reservoir tube was mixed with TRIzol LS reagent and processed for RNA purification.

For EDTA pretreatment, the lysate was treated with 10 mM EDTA for 30 min on ice and then mixed with an additional 10 mM MgCl_2_ before RNase I digestion. Denaturing UREA-PAGE gel was stained with SYBR Gold (Thermo Fisher Scientific).

For rRNA subtraction, stand-alone Ribo-Zero kits [Ribo-Zero Gold rRNA Removal Kit (Human/Mouse/Rat)] (Illumina), Ribo-Zero accompanied by a TruSeq Stranded Total RNA Kit (Illumina), and riboPOOL kits (*Drosophila melanogaster*, *Saccharomyces cerevisiae*, pan-plant, and pan-bacteria; siTOOLs Biotech) were used.

#### Harringtonine chase assay

Cells were treated with 2 μg/ml harringtonine (MedChemExpress) at 37°C for 60, 120, or 180 s. Then, the cells were treated with 100 μg/ml cycloheximide and incubated for 1 min at 37°C before cell harvesting. For no-chase samples, harringtonine treatment was omitted, but cycloheximide was added to the medium.

#### Library preparation for RNA-Seq

For RNA-Seq library preparation, 0.5 µg of total RNA (with a TruSeq Stranded mRNA Library Prep Kit; Illumina) or 5 ng of total RNA (with a SMARTer Stranded Total RNA-Seq Kit v3; Takara Bio) was used.

#### Sequencing

Libraries were sequenced on a HiSeq 4000 or HiSeq X system (Illumina) with the paired-end/150 nt-long read option.

#### Data analysis

Sequence data were processed as previously described ^96^. The changes in the read counts on the ORFs/transcripts were calculated with the DESeq2 package ^97^ and then renormalized to the mean change in mitochondrial mRNAs (as internal spike-ins) or Rluc and Fluc mRNAs (as external spike-ins) to measure overall alteration.

The absolute ribosome number (*N*) on transcript *j* was determined as follows:

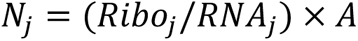

where *Ribo_j_* represents ribosome footprint reads per million mapped reads (RPM) on the ORF of transcript *j,* and *RNA_j_* represents the transcript per million (TPM) of transcript *j*. The coefficient *A* was defined as follows:

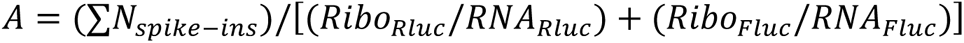

Theoretically, the *Ribo_Rluc_/RNA_Rluc_* and *Ribo_Fluc_/RNA_Fluc_* ratios are 2 and 3, respectively. Thus, the sum of the ribosome number (∑*N_spike-ins_*) should be 5. Transcripts with RPM ≥ 1 and TPM ≥ 1 were considered.

The absolute ribosome number (*M*) on each codon *k* of transcript *j* was determined as follows:

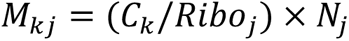

where *C_k_* represents the ribosome footprint RPM on codon *k*.

For the ribosome run-off assay, smoothed metagene profiles were obtained from the transcripts possessing an ORF with 2000 or more codons. The mean read density from 1000 to 1500 codons was set to 1.

The translation initiation rate (*I*) for transcript *j* was defined as follows:

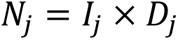

where *D_j_* represents the duration of the loaded ribosome on transcript *j*. Given that the loaded ribosome undergoes a translation elongation reaction on the ORF of length *L_j_* in transcript *j*, *D_j_* was defined as follows:

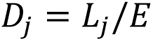

where *E* represents the global elongation rate, which was determined to be 4.06 codon/s by a ribosome run-off assay. Thus,

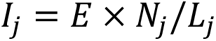

The lifetime translation round (*R*) for transcript *j* was defined as follows:

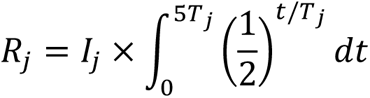

where *T_j_* represents the half-life of transcript *j*, as determined by BRIC-Seq. Here, we calculated the ribosome loading frequency during 5*T_j_*, when ∼97% of the mRNA was predicted to be degraded.

> All the data are presented as the average of replicates when available.

> Disome footprint data (GSE145723) ^19^ were processed as described above.

> The TOP mRNAs (n = 614) were defined as described in an earlier study ^98^. The mRNAs possessing uORFs (n = 21) were identified in a previous study ^25^.

### BRIC-Seq

#### Library preparation

The libraries prepared in this study are summarized in Supplementary Data 4. BRIC-Seq was performed as described previously, with modifications ^56^. HEK293T cells expressing control shRNA ^19^ were seeded in 10-cm dishes and incubated in 10 ml of DMEM supplemented with 150 µM bromouridine (BrU; FUJIFILM Wako Pure Chemical Corporation) for 24 h. The cells were then washed twice with 5 ml of DMEM without BrU, incubated in 10 ml of DMEM without BrU for 0, 2, 4, 6, 10, or 24 h, washed with 5 ml of PBS, and lysed by adding 600 µl of TRIzol reagent (Thermo Fisher Scientific) directly to the dishes. Total RNA was purified using a Direct-zol RNA MiniPrep Plus Kit (Zymo Research).

BrU-containing spike-in mRNAs were synthesized via *in vitro* transcription. The template DNA fragments were PCR-amplified using the psiCHECK2 plasmid (Promega) with 5′-TAATACGACTCACTATAGG-3′ and 5′-CACACAAAAAACCAACACACAG-3′ (Rluc); the psiCHECK2 plasmid with 5′-ACTTAATACGACTCACTATAGGAAGCTTGGCATTCCGG-3′ and 5′-TGTATCTTATCATGTCTGCTCGAAG-3′ (Fluc); and the pColdI-GFP plasmid ^99^ with 5′-TGACTAATACGACTCACTATAGGATCTGTAAAGCACGCCATATCG-3′ and 5′-TGGCAGGGATCTTAGATTCTG-3′ (GFP). *In vitro* transcription was performed using a T7-Scribe Standard RNA IVT Kit (CELLSCRIPT) in the presence of 1.2 mM BrUTP (Jena Bioscience) and 5 mM UTP. The RNA was purified with RNAClean XP beads (Beckman Coulter).

For immunoprecipitation, 25 µl of Dynabeads M-280 Sheep Anti-Mouse IgG (Thermo Fisher Scientific) was incubated with 2 mg of anti-BrdU antibody (BD Biosciences, 555627) for 3 h at 4°C and equilibrated with BRIC-Seq lysis buffer (lysis buffer without cycloheximide and chloramphenicol). Twenty micrograms of total RNA mixed with a spike-in RNA mixture (1 ng of Rluc, 0.2 ng of Fluc, and 0.04 ng of GFP) was incubated with the magnetic beads in BRIC-Seq lysis buffer for 2 h at 4°C, washed 5 times with 100 µl of BRIC-Seq lysis buffer, and mixed with 200 µl of TRIzol reagent. RNA was extracted with a Direct-zol RNA MicroPrep Kit and subjected to RNA-Seq library preparation using a Ribo-Zero Gold rRNA Removal Kit (Human/Mouse/Rat) (Illumina) followed by a TruSeq Stranded Total RNA Kit (Illumina).

#### Sequencing

Libraries were sequenced on a HiSeq 4000 (Illumina) with the single-end/50 nt-long read option.

#### Data analysis

Filtering, trimming of adapters, and mapping to the genome were performed as described in the RNA-Seq section. The fold changes in the read counts compared to those of the 0 h samples were calculated with the DESeq2 package ^97^ and then renormalized to the mean change resulting from three spike-ins. For the calculation of the mRNA half-life, the log_2_-converted fold change at each time point was fitted to a linear model. Since the decay of transcripts sometimes slows at later time points, we calculated the half-life using all combinations of earlier samples (*i.e.*, 0-4 h, 0-6 h, 0-10 h, and 0-24 h) and fitted the results with the maximum R^2^ value. Transcripts with R^2^ > 0.9 and 0 < half-life < 24 h were used in downstream analysis.

### Polysome profiling and RT‒qPCR

HEK293 T-REx cells cultured in a 10-cm dish were lysed in 600 μl of SDG buffer (20 mM Tris-HCl pH 7.5, 150 mM NaCl, 5 mM MgCl_2_, 1% Triton X-100, 1 mM DTT, and 100 µg/ml cycloheximide). Two hundred microliters of the lysate were loaded onto a 10-50% sucrose gradient and separated by ultracentrifugation as described in the “*Preparation of spike-in Rluc-disome and Fluc-trisome*” section above. The total RNA in each fraction and input lysate was extracted using TRIzol LS reagent and a Direct-zol RNA MicroPrep Kit. cDNAs were synthesized using the ReverTraAce qPCR RT Kit (TOYOBO), according to the manufacturer’s protocols. RT‒qPCR analyses were conducted using an iTaq Universal SYBR Green Supermix Kit (Bio-Rad) and a CFX Connect Real Time System (Bio-Rad). Three independently passaged biological replicates were analyzed. The following primers were utilized for the qRT‒PCR assays: *MED29*, 5′-TCCTCAGCTGCTGGTGTATCG-3′ and 5′-TCACTGCTCTTTCTGTAGACTCTCC-3′; *RPL27*, 5′-ATGGGCAAGTTCATGAAACCTGGGA-3′ and 5′-TTATTGACGACAGTTTTGTCCAAGG-3′; *PCLAF*, 5′-CAGAAAAGTGGTGGCTGCTCGAGCC-3′ and 5′-CAAAGGACATGCTCTTTCCTCGATG-3′; and *PET100*, 5′-TCACTTTCCCTGTGGCTATGTTCTG-3′ and 5′-TCATGGCTTCTCAGGTGGCCACAGC-3′.

### Nascent peptide labeling with OP-puro

Cells seeded in 6-well plates were treated with 20 μM OP-puro (Jena Bioscience) at 37°C for 30 min, washed with 1 ml of phosphate-buffered saline (PBS), and lysed in 100 µl of OP-puro lysis buffer (20 mM Tris-HCl pH 7.5, 150 mM NaCl, 5 mM MgCl_2_, and 1% Triton X-100). The lysate was then incubated with 1 μM IRDye 800CW azide (LI-COR Bioscience) for 30 min at 25°C using a Click-iT Cell Reaction Buffer Kit (Thermo Fisher Scientific). After the removal of unreacted azide by MicroSpin G-25 columns (Cytiva), the proteins were separated by sodium dodecyl sulfate (SDS)-polyacrylamide gel electrophoresis (PAGE). The infrared (IR) 800 signal was then detected using an ODYSSEY CLx (LI-COR Biosciences). The total protein on the gel was stained with EzStain AQua (ATTO) and detected by ODYSSEY CLx (LI-COR Biosciences) with an IR700 channel. Total protein-normalized nascent peptide signals were quantified after subtraction of the background.

### Western blotting

The following primary antibodies were used: anti-HSP70 family proteins (Cell Signaling Technology, #4872; 1:1000) and anti-β-actin (LI-COR Biosciences, #926-42212; 1:1000). For the secondary antibodies, we utilized IRDye 800CW anti-rabbit IgG (LI-COR Biosciences, #926-32211: 1:10000) and IRDye 680RD anti-mouse IgG (LI-COR Biosciences, #925-68070: 1:10000). The images were subsequently captured using an ODYSSEY CLx system (LI-COR Biosciences).

### Cell viability assay

Cells were seeded in white 96-well plates and incubated with the RealTime-Glo MT Cell Viability Assay (Promega). Luminescence was detected using a GloMax-96 instrument (Promega).

## Data Availability

The ribosome profiling and RNA-Seq data obtained in this study (GSE233555) were deposited in the National Center for Biotechnology Information (NCBI) database. This study also used deposited disome profiling data (GSE145723) ^19^ and BRIC-Seq data (GSE233374). Source data are provided with this paper.

## Code availability

The key custom scripts used in this study are available at Zenodo (https://zenodo.org, DOI: 10.5281/zenodo.10223916).

## Acknowledgments

We are grateful to all the members of the Iwasaki laboratory for their constructive discussions and technical help. We thank Dr. Yuichiro Mishima for his critical comments. We are grateful to Dr. Yusuke Kimura, Dr. Tomoya Fujita, and Dr. Shiho Makino for sharing the lysates. Computation was supported by the HOKUSAI SailingShip supercomputer facility at RIKEN. This work was supported by the Ministry of Education, Culture, Sports, Science and Technology (MEXT) [a Grant-in-Aid for Transformative Research Areas (B) “Parametric Translation”, JP20H05784 (to S.I.), JP20H05786 (to Y.I.), and JP20H05782 (to M.D.); a Grant-in-Aid for Transformative Research Areas (A) “Multifaceted Proteins”, JP21H05734 and JP23H04268 (to Y.S.)], the Japan Agency for Medical Research and Development (AMED) [AMED-CREST, JP20gm1410001 (to S.I. and Y.I.); AMED-PRIME, JP23gm6910005 (to Y.S.); Research Program on Emerging and Re-emerging Infectious Diseases, JP22fk0108570 (to H.T.)], the Japan Society for the Promotion of Science (JSPS) [a Grant-in-Aid for Scientific Research (B), JP23H02415 (to S.I.); a Grant-in-Aid for Early Career Scientists, JP21K15023 (to Y.S.); a Grant-in-Aid for Scientific Research (C), JP23K05648 (to Y.S.); a Grant-in-Aid for Research Activity Start-up, JP22K20765 (to H.T.); a Grant-in-Aid for Early-Career Scientists, JP23K14173 (to H.T.); a Grant-in-Aid for JSPS Fellows, JP23KJ2178 (to H.T.) and JP23KJ2175 (to N.K.)], the Japan Science and Technology Agency (JST) [FOREST Program, JPMJFR226F (to T.M.)], and RIKEN [Pioneering Projects “Biology of Intracellular Environments” (to S.I. and Y.S.)]. K.T. was supported by a RIKEN Student Researcher program and a World-leading Innovative Graduate Study Program in Proactive Environmental Studies (WINGS-PES) from The University of Tokyo. Part of the next-generation sequencing was conducted by HiSeq 4000 [supported by the National Institutes for Health (NIH) Instrumentation Grant (S10 OD018174)] in QB3 Genomics, UC Berkeley, Berkeley, CA, (RRID:SCR_022170). H.T. and N.K. were the JSPS Research Fellows (PD).

## Author contributions

Conceptualization, K.T., M.M., Y.S., and S.I.; Methodology, K.T., M.M., Y.S., and S.I.; Formal analysis, K.T., M.M., H.T., N.K., T.M., S.Y.A.C., and Y.S.; Investigation, K.T., M.M., H.T., N.K., T.M., S.Y.A.C., and Y.S.; Writing – Original Draft, K.T., Y.S., and S.I.; Writing – Review & Editing, K.T., M.M., H.T., N.K., T.M., S.Y.A.C., M.D., Y.I., Y.S., and S.I.; Visualization, K.T.; Supervision, M.D., Y.I., Y.S., and S.I.; Project administration, S.I.; Funding Acquisition, H.T., N.K., T.M., M.D., Y.I., Y.S., and S.I.

---

**Supplementary Fig. 1:**
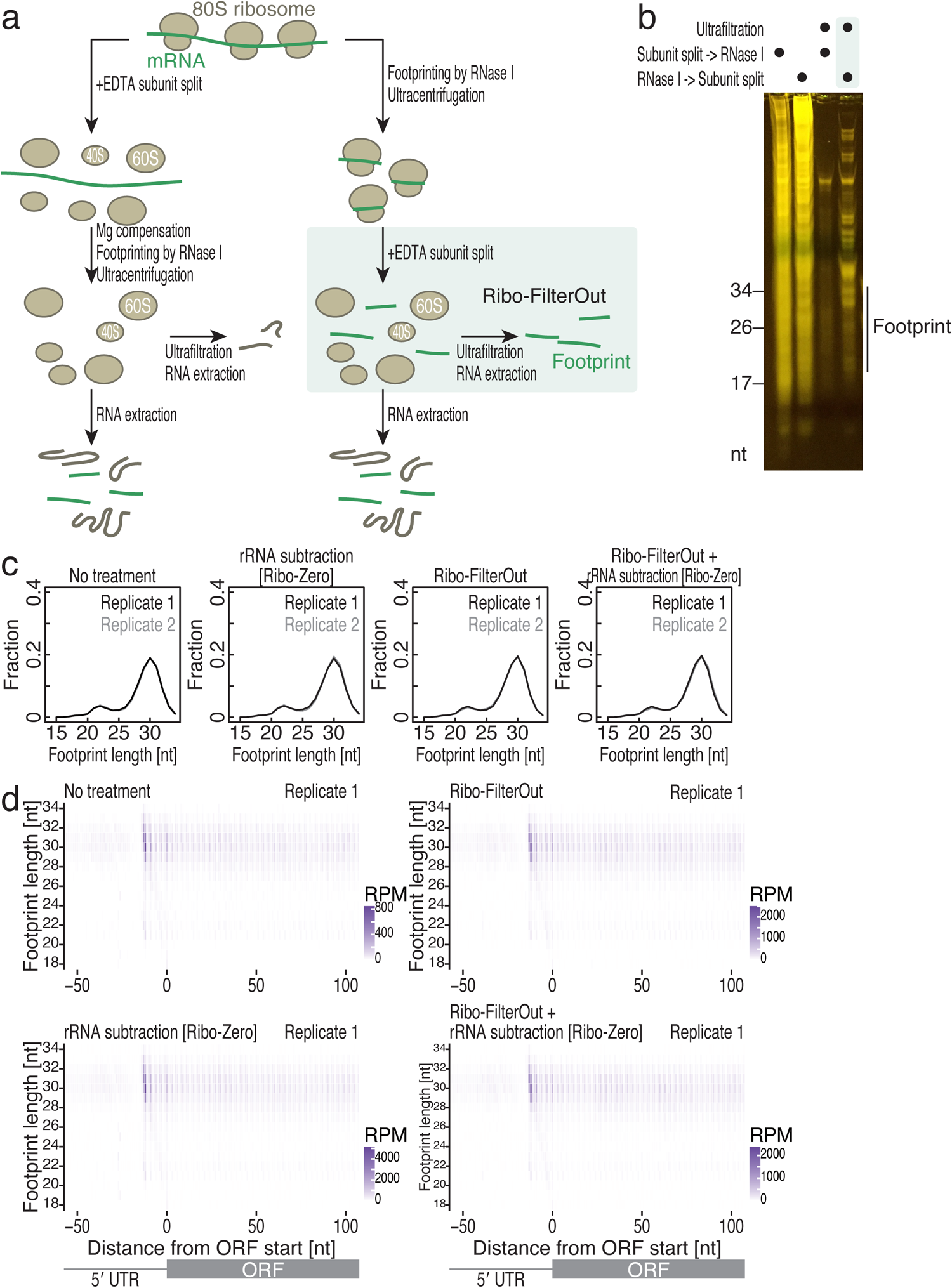
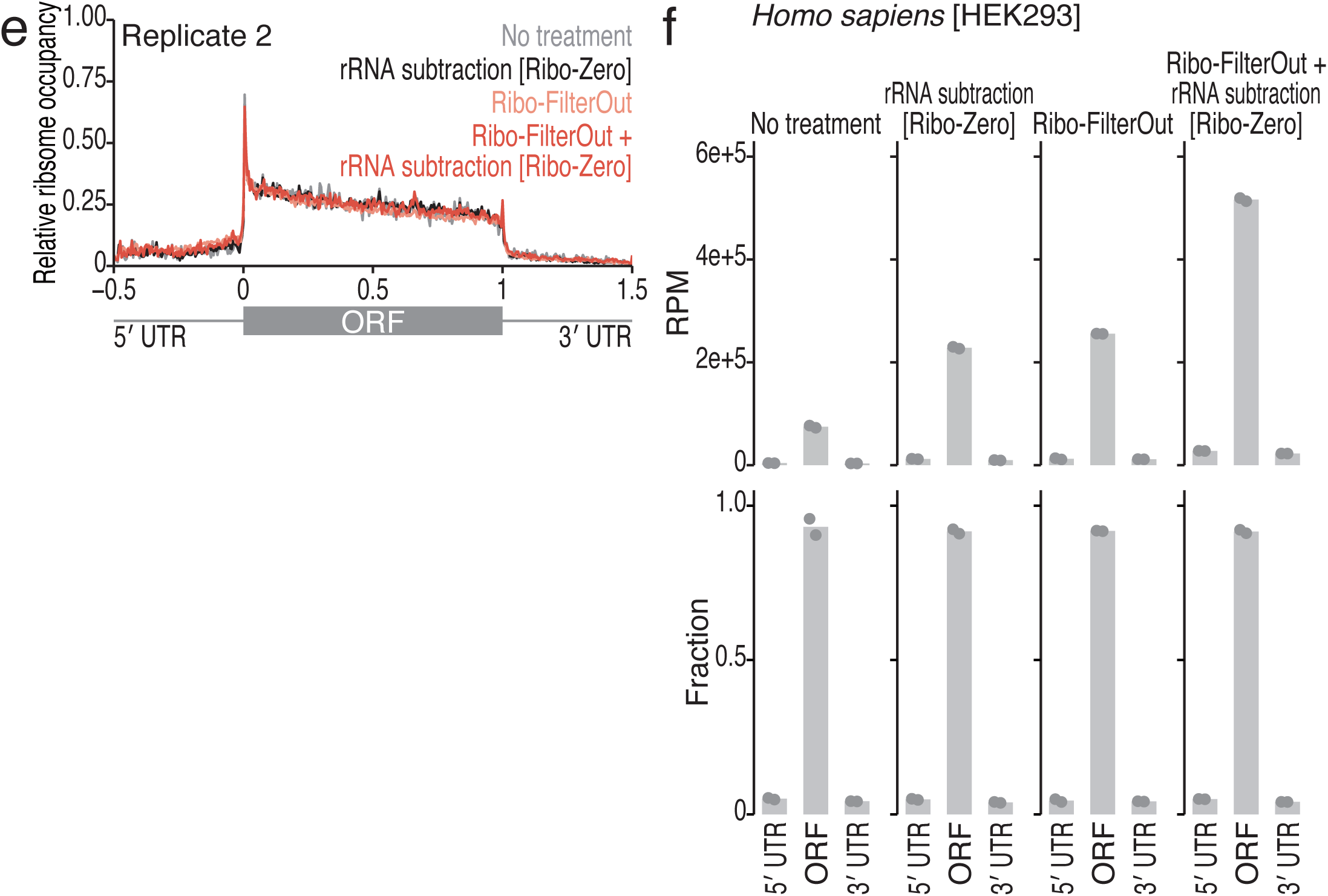
Characterization of the Ribo-FilterOut technique. (a) Schematic of the experiments in b. Right: mRNA-ribosome complexes were treated with RNase I, after which the ribosomal complex was isolated via ultracentrifugation. Ribosome footprints were separated from ribosome subunits by EDTA treatment and then isolated by ultrafiltration. Left: For the control, the lysate was pretreated with EDTA to release mRNA from ribosomes before RNase I digestion. (b) SYBR Gold staining of the RNA fragments obtained from the experiments shown in a. (c) The distribution of ribosome footprint length in ribosome profiling for the indicated conditions. (d) Metagene plots of the 5′ end of ribosome footprints around the start codon (the first position of the start codon was set to 0) along the footprint length. The color scales for read abundance are shown. (e) Metagene plot of the relative ribosome footprint distribution (ribosome occupancy) along the 5′ UTR, ORF, and 3′ UTR. The length of the ORF was set to 1, whereas the lengths of the 5′ UTR and 3′ UTR were set to 0.5. (f) Reads obtained from ribosome profiling were mapped to the indicated region of mRNAs (top). The relative distributions of the 5′ UTR, ORF, and 3′ UTR are shown (bottom). The mean (bar) and individual replicates (n = 2, points) are shown. RPM, reads per million mapped reads.

**Supplementary Fig. 2:**
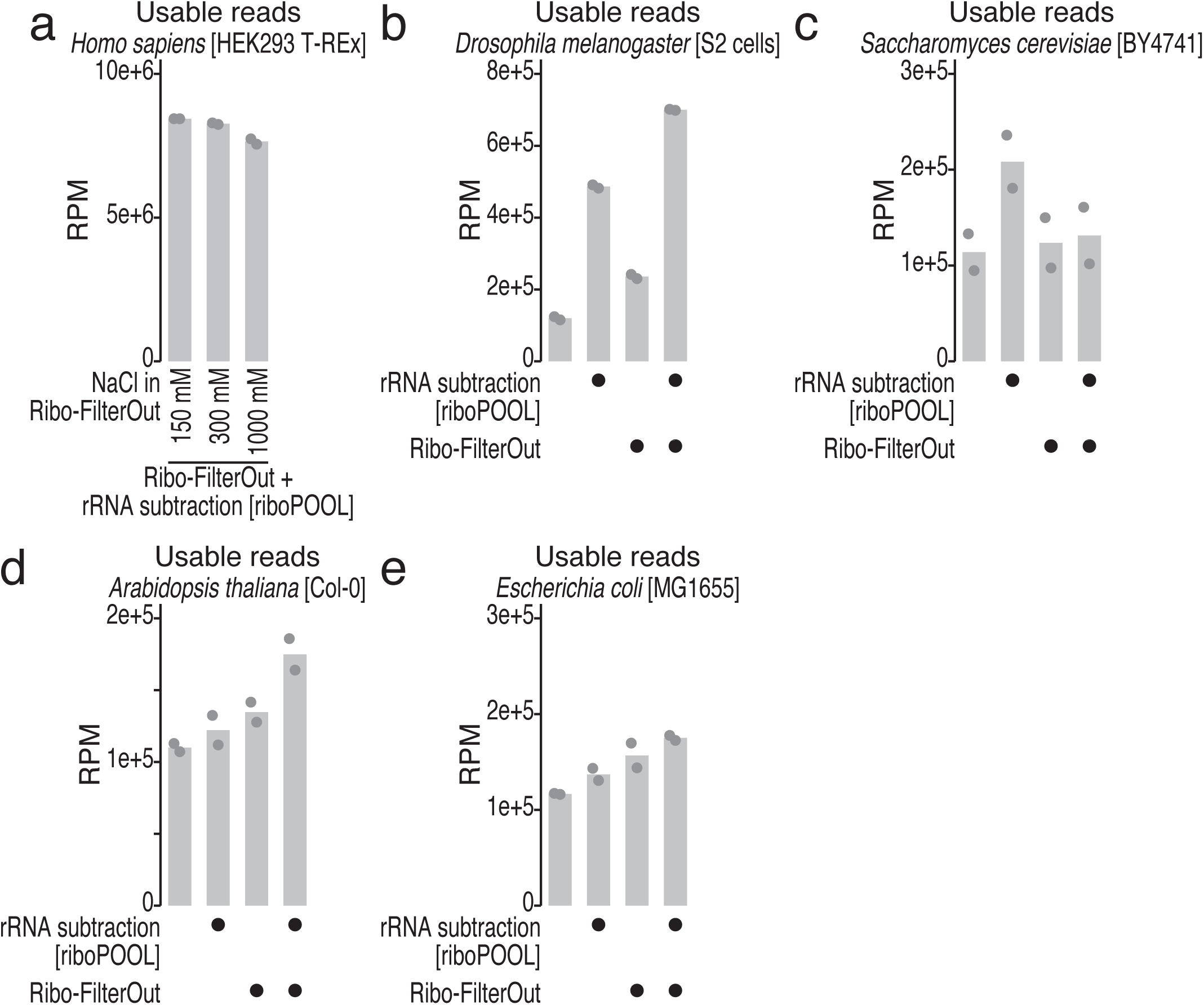
Application of Ribo-FilterOut to various species. (a) Test of salt concentrations in the resuspension buffer of the ribosome pellet after sucrose cushion centrifugation. The amount of usable reads is shown. (b-e) Combination of the standard rRNA subtraction method (riboPOOL) with Ribo-FilterOut for fly S2 cells (b), yeast (c), plants (d), and bacteria (e). The amount of usable reads is shown. The mean (bar) and individual replicates (n = 2, points) are shown. RPM, reads per million mapped reads.

**Supplementary Fig. 3:**
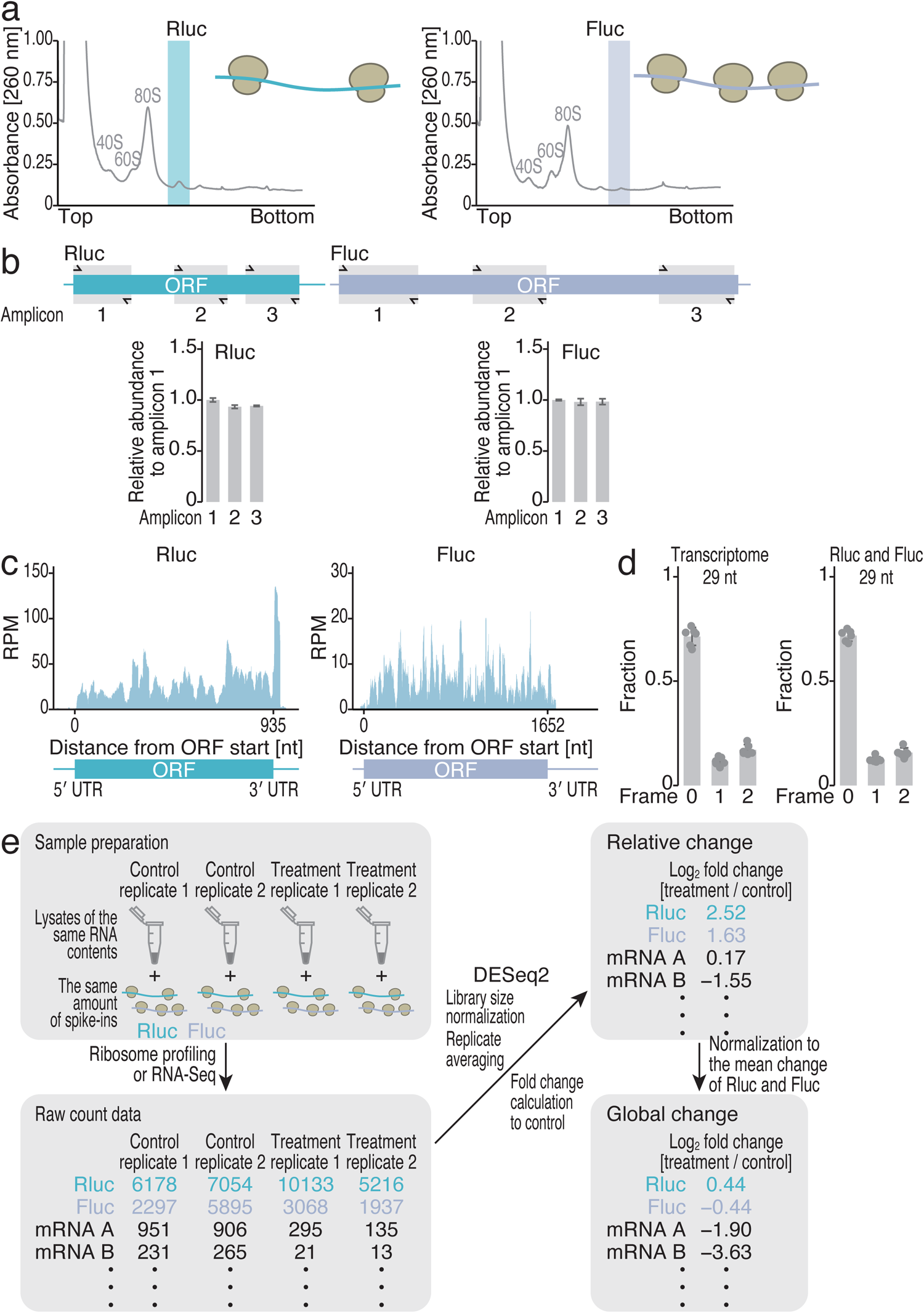
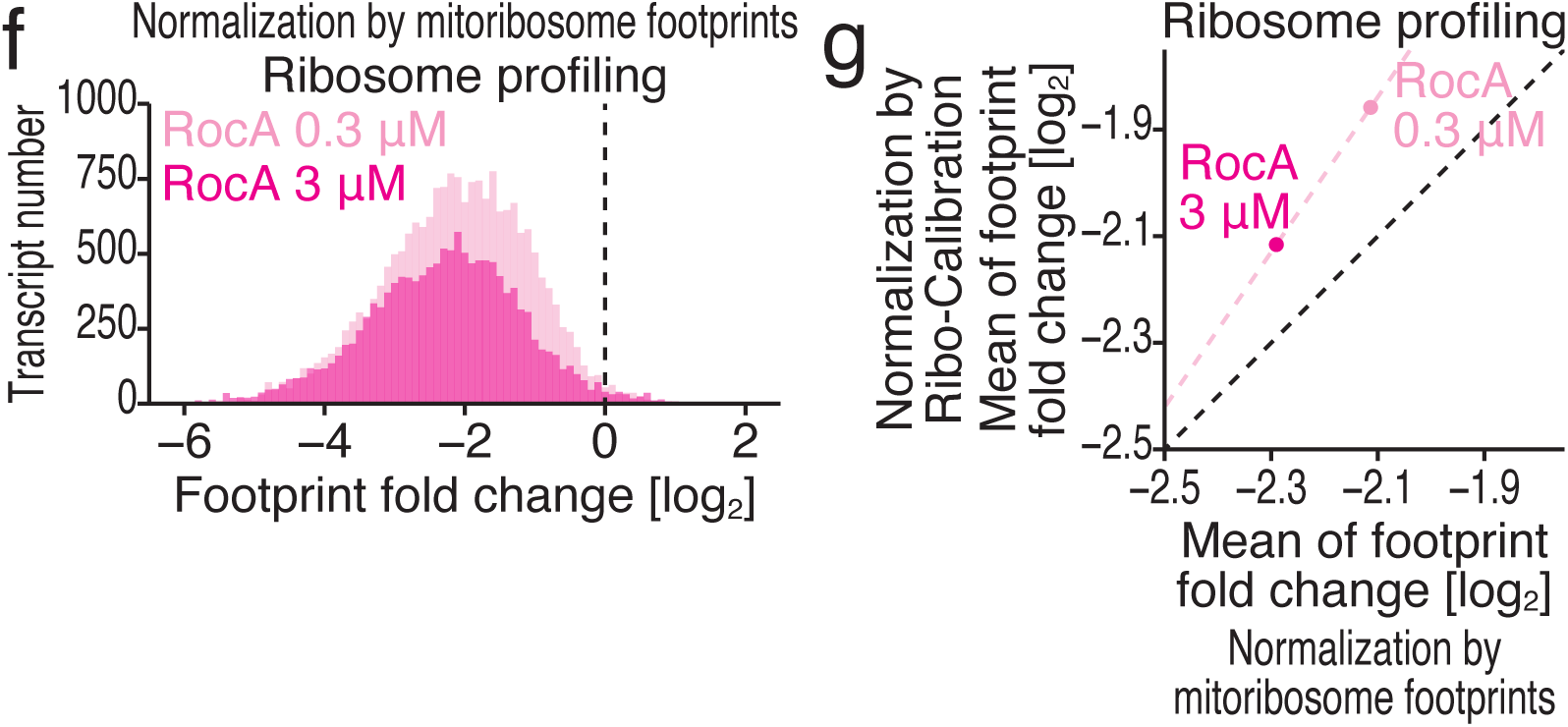
Characterization of the method for global translation measurement by Ribo-Calibration. (a) Sucrose density gradient ultracentrifugation for the separation of ribosomal complexes. Rluc mRNA (left) and Fluc mRNA (right) were translated *in vitro* with RRL. Highlighted fractions were isolated for spike-ins for ribosome profiling. (b) RT‒qPCR analyses of Rluc and Fluc mRNAs complexed with ribosomes. Three different amplicons were designed for each transcript (top). The data were calibrated with titrated amounts of bare mRNAs used for *in vitro* translation as standards. The mean (bar) of three replicates and s.d. (error) are shown. (d) Ribosome footprint coverage on RLuc mRNA (left) and Fluc mRNA (right). RPM, reads per million mapped reads. (d) Fraction of the frame position where the 5′ ends of the reads are located. Twenty-nine nucleotide footprints were analyzed. The mean (bar), individual data (n = 6; 2 replicates for DMSO, 2 replicates for 0.3 μM RocA treatment, and 2 replicates for 3 μM RocA treatment, points), and s.d. (error) are shown. (e) Schematic of sample handling and data calculation for the measurement of global translation changes. The use of the same amount of lysate and spike-ins of mRNA-ribosome complexes was essential. (f) Histograms of ribosome footprint fold changes after 0.3 or 3 μM RocA treatment normalized to the mitochondrial footprint. (g) Comparison of the changes in global translation caused by RocA treatment calculated from Ribo-calibration and mitoribosome footprints.

**Supplementary Fig. 4:**
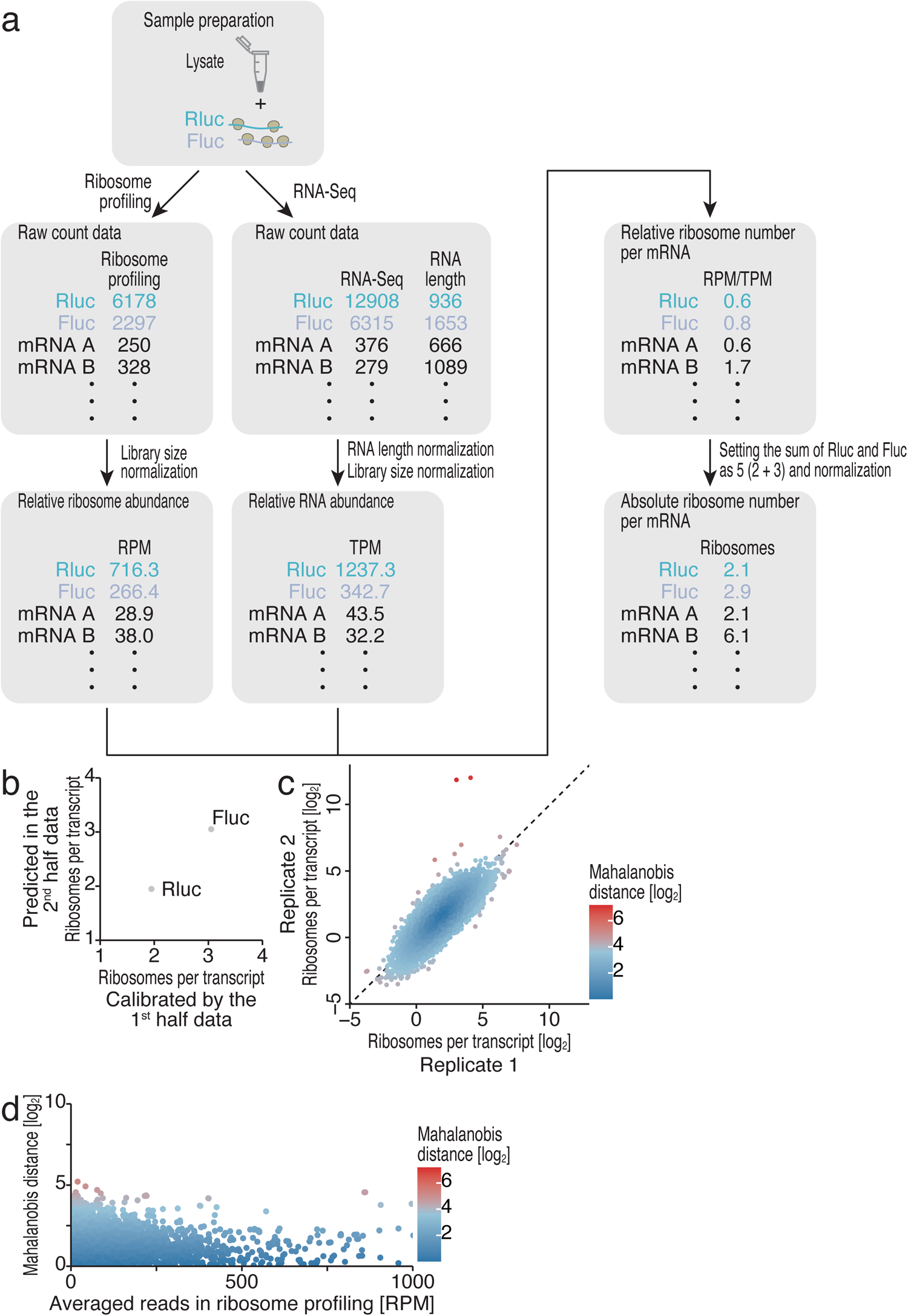
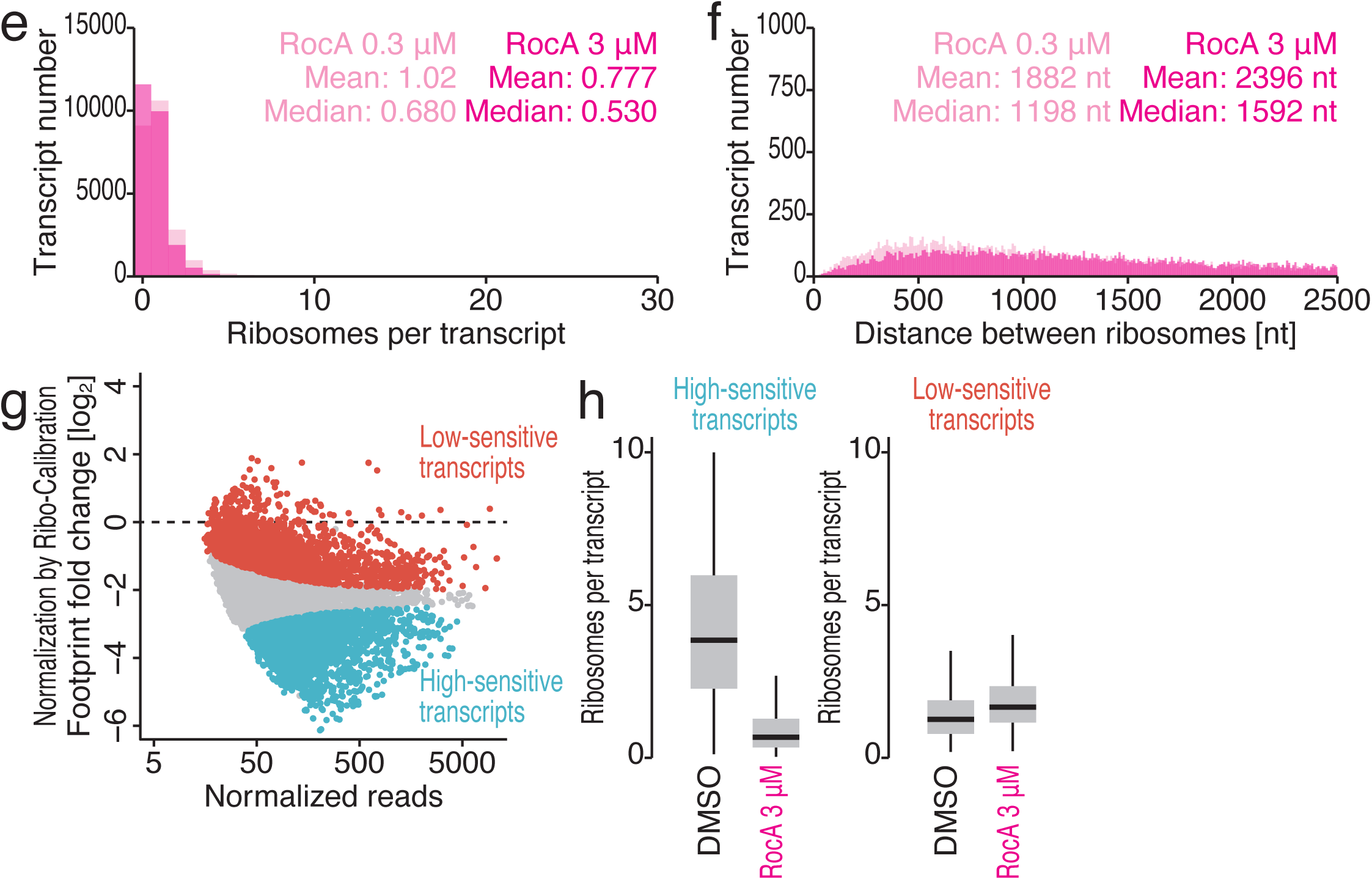
Characterization of the method for the absolute calibration of ribosome number by Ribo-Calibration. (a) Schematic of sample handling and data calculation for the absolute quantification of ribosome numbers on ORFs. Ribosome profiling and RNA-Seq must be performed using the same lysate supplemented with purified mRNA-ribosome complexes. (b) To validate the performance of Ribo-Calibration for the absolute quantification of ribosome numbers on ORFs, we divided the data into halves, used the first half for calibration of the data, and then predicted the ribosome numbers on Rluc and Fluc in the second half. (c) Scatter plot of ribosome numbers on ORFs in replicates, scoring the reproducibility with the Mahalanobis distance (shown in the color scale). (d) Scatter plot of the Mahalanobis distance (shown in c) and average ribosome footprint reads from replicates. The color scales for the Mahalanobis distance are shown. RPM, reads per million mapped reads. (e) Histograms of the absolute ribosome number on transcripts with 0.3 or 3 μM RocA treatment. (f) Histograms of the distance between ribosomes with 0.3 or 3 μM RocA treatment. (g) MA (M, log ratio; A, mean average) plot of the ribosome footprint fold change with 3 μM RocA treatment normalized by Ribo-Calibration spike-ins. Transcripts with significant changes (FDR < 0.05) are highlighted. (h) Box plots of the absolute number of ribosomes for RocA high-sensitivity transcripts (left) and low-sensitivity transcripts (right). In box plots, the median (centerline), upper/lower quartiles (box limits), and 1.5× interquartile range (whiskers) are shown.

**Supplementary Fig. 5:**
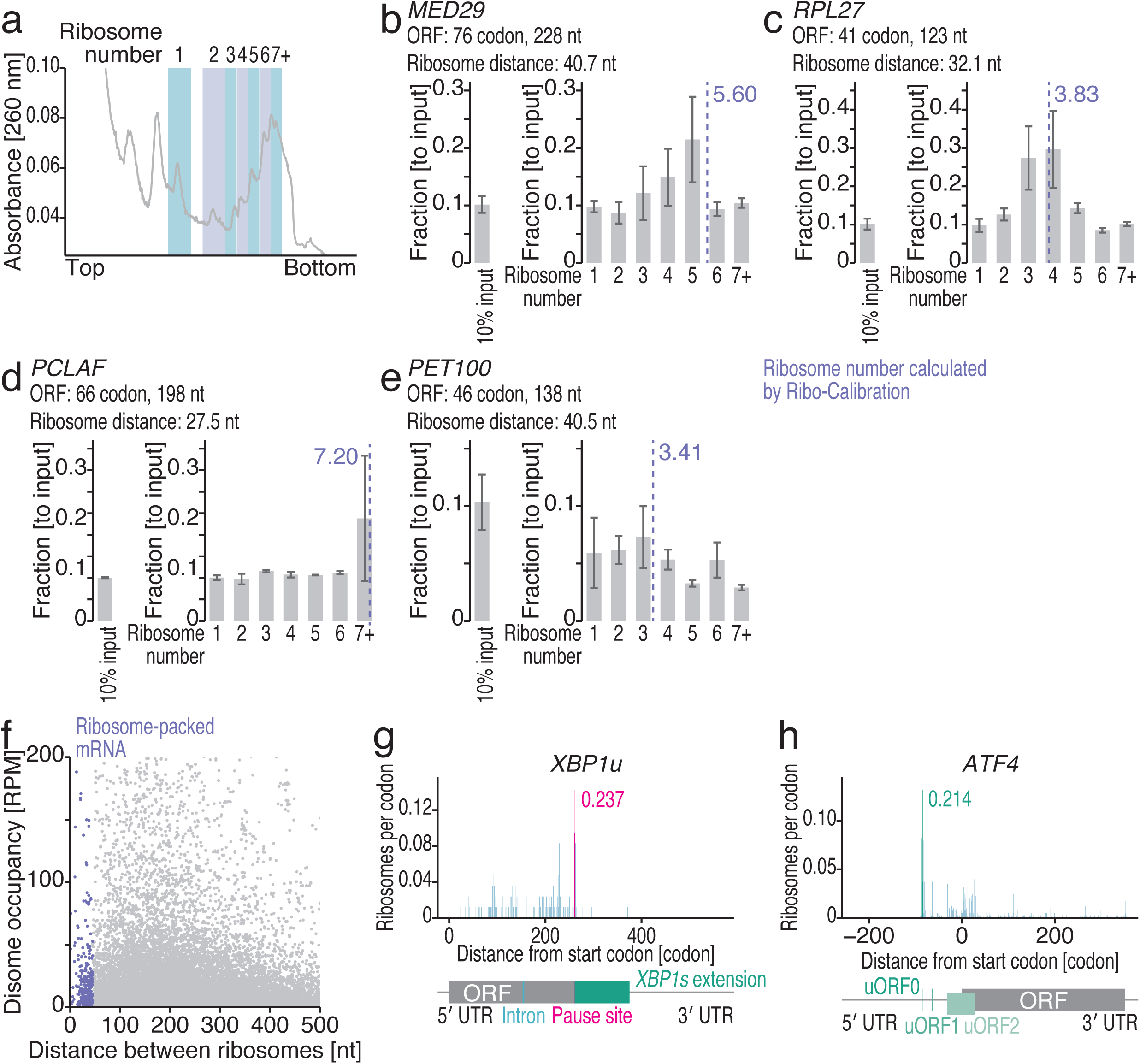
Validation of ribosome numbers on ORFs obtained by Ribo-Calibration. (a) Sucrose density gradient ultracentrifugation for the separation of ribosomal complexes from HEK293 T-REx cells, highlighting the fractions used for downstream RT‒qPCR analyses. (b-e) RT‒qPCR analysis of the indicated mRNAs in the polysome fractions according to the number of ribosomes on the transcripts. Ten percent of the input lysate was used for the standard. The mean (bar) of three replicates and s.d. (error) are shown. (f) Scatter plots of the distance between ribosomes and disome occupancy. RPM, reads per million mapped reads. (g and h) Distribution of ribosome footprints along the indicated genes. The A-site position of the reads is depicted. The calibrated ribosome numbers on the pause site in *XBP1u* and uORF0 in *ATF4* (including ribosomes on codons before and ahead) are highlighted.

**Supplementary Fig. 6:**
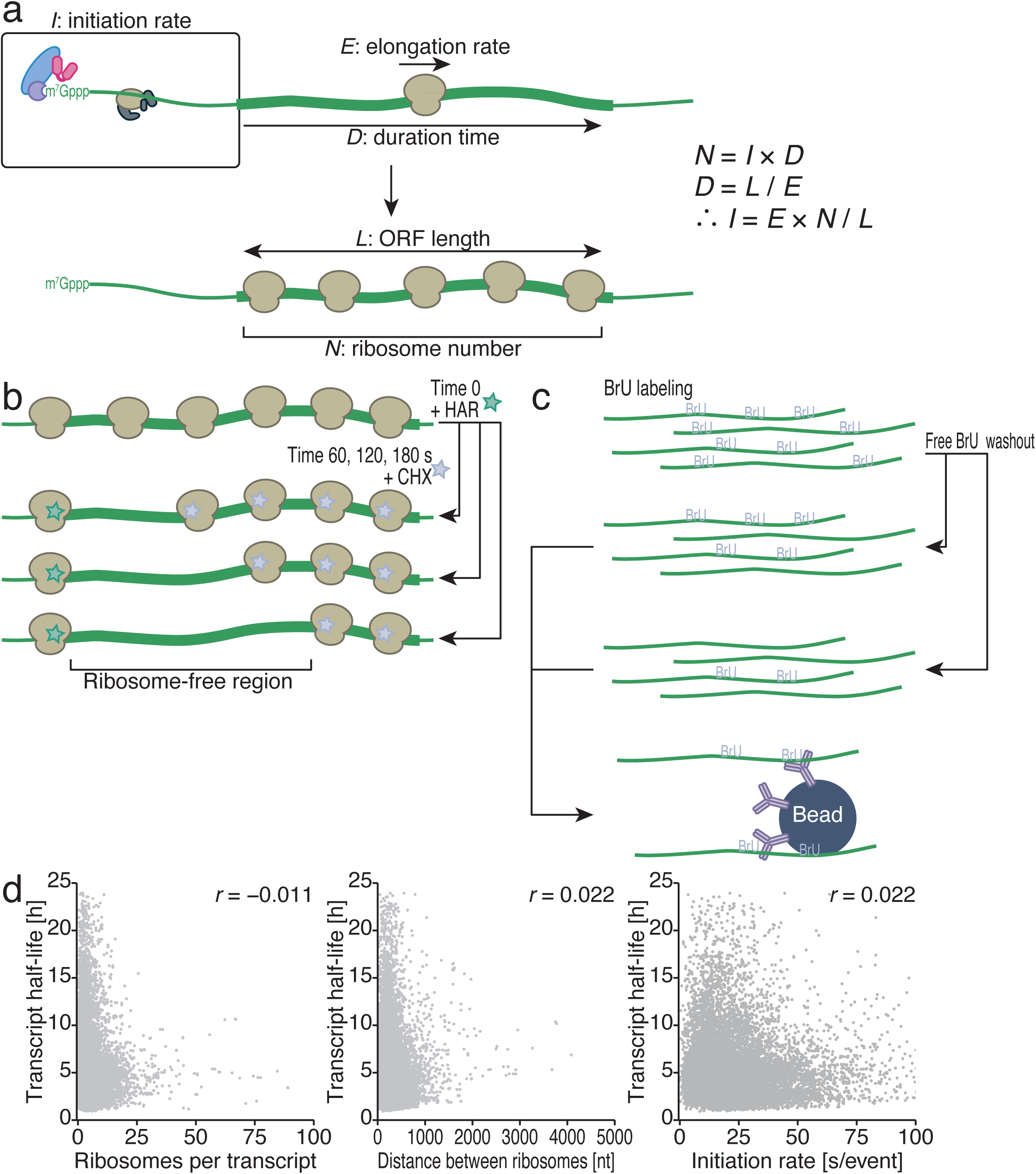
Characterization of ribosome run-off assay and BRIC-Seq. (a) Schematic representation of the relationships among the number of ribosomes on the ORF (*N*), the duration of ribosome residence on the ORF (*D*), the length of the ORF (*L*), the translation elongation rate (*E*), and the translation initiation rate (*I*). (b) Schematic of the ribosome run-off assay with harringtonine (HAR) chase. After a short incubation with HAR, ribosomes on the transcripts were fixed with cycloheximide (CHX). (c) Schematic of BRIC-Seq with bromouridine (BrU). After the labeling of cellular mRNA with BrU, the cells were incubated in medium without BrU over the time course. mRNAs were isolated with an anti-BrdU antibody. (d) Comparison of transcript half-life and the absolute number of ribosomes on transcripts (left), the distance between ribosomes (middle), and the translation initiation rate (right). *r*, Pearson’s correlation coefficient.

**Supplementary Fig. 7:**
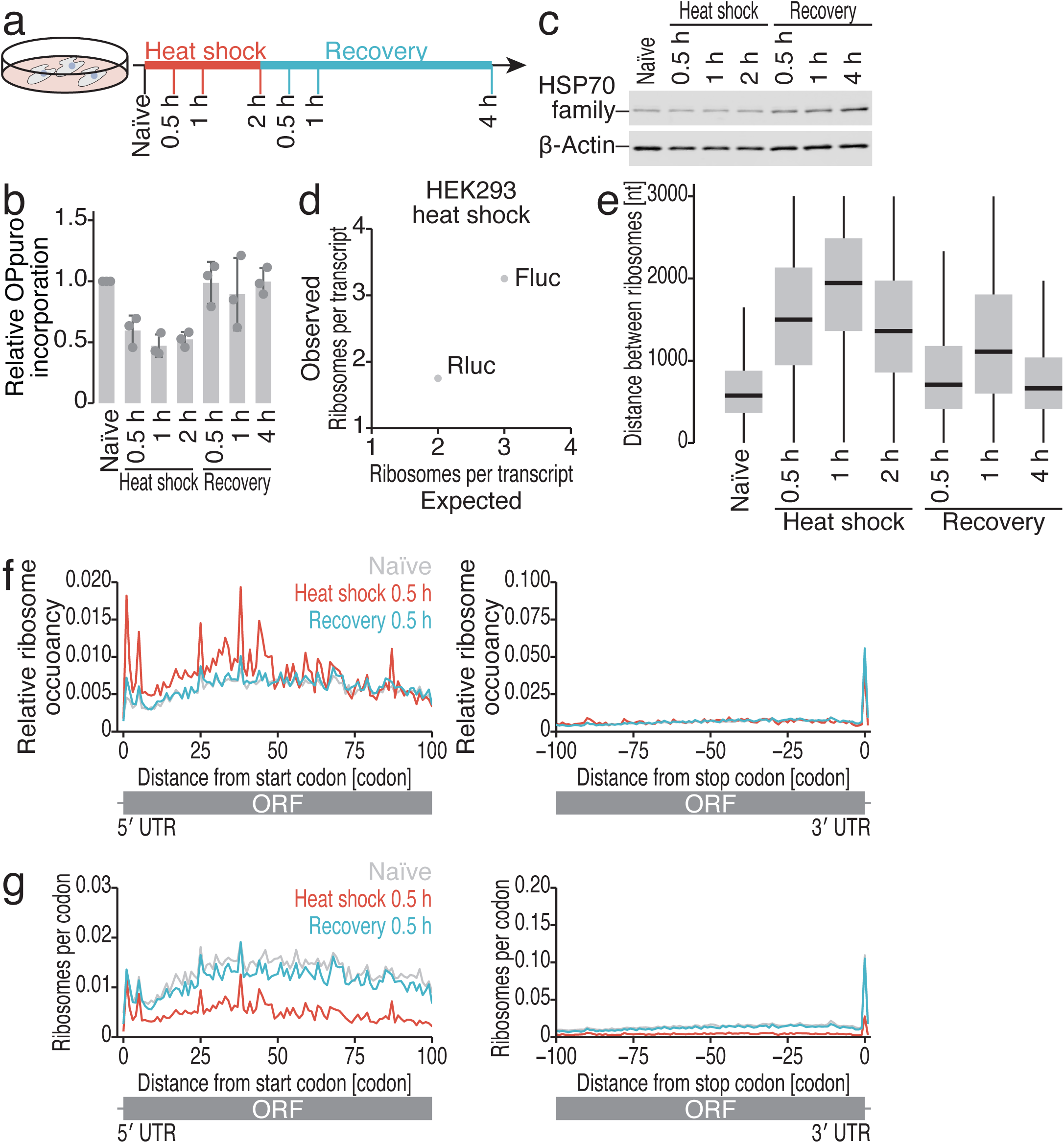
Characterization of heat shock and recovery in HEK293 cells. (a) Schematic of the experiments over the time course. (b) Relative OP-puro incorporation during heat shock and recovery. The mean (bar), s.d. (error), and individual data (n = 3, points) are shown. (c) Western blotting of the indicated proteins. The anti-HSP70 antibody used in this study detected both heat-inducible HSP70 family proteins and constitutively expressed HSP70 (Hsc70) family proteins. (d) Comparison of expected and observed ribosome numbers on Rluc and Fluc transcripts used for Ribo-Calibration. (e) Box plots of the distance between ribosomes during heat shock and recovery. (f) Metagene plots with relative ribosome occupancy at each position around start codons (left, the first position of the start codon was set to 0) and stop codons (right, the first position of the stop codon was set to 0) under the indicated conditions. (g) Metagene plots with calibrated ribosome numbers at each position around start codons (left, the first position of the start codon was set to 0) and stop codons (right, the first position of the stop codon was set to 0) under the indicated conditions. In box plots, the median (centerline), upper/lower quartiles (box limits), and 1.5× interquartile range (whiskers) are shown.

**Supplementary Fig. 8:**
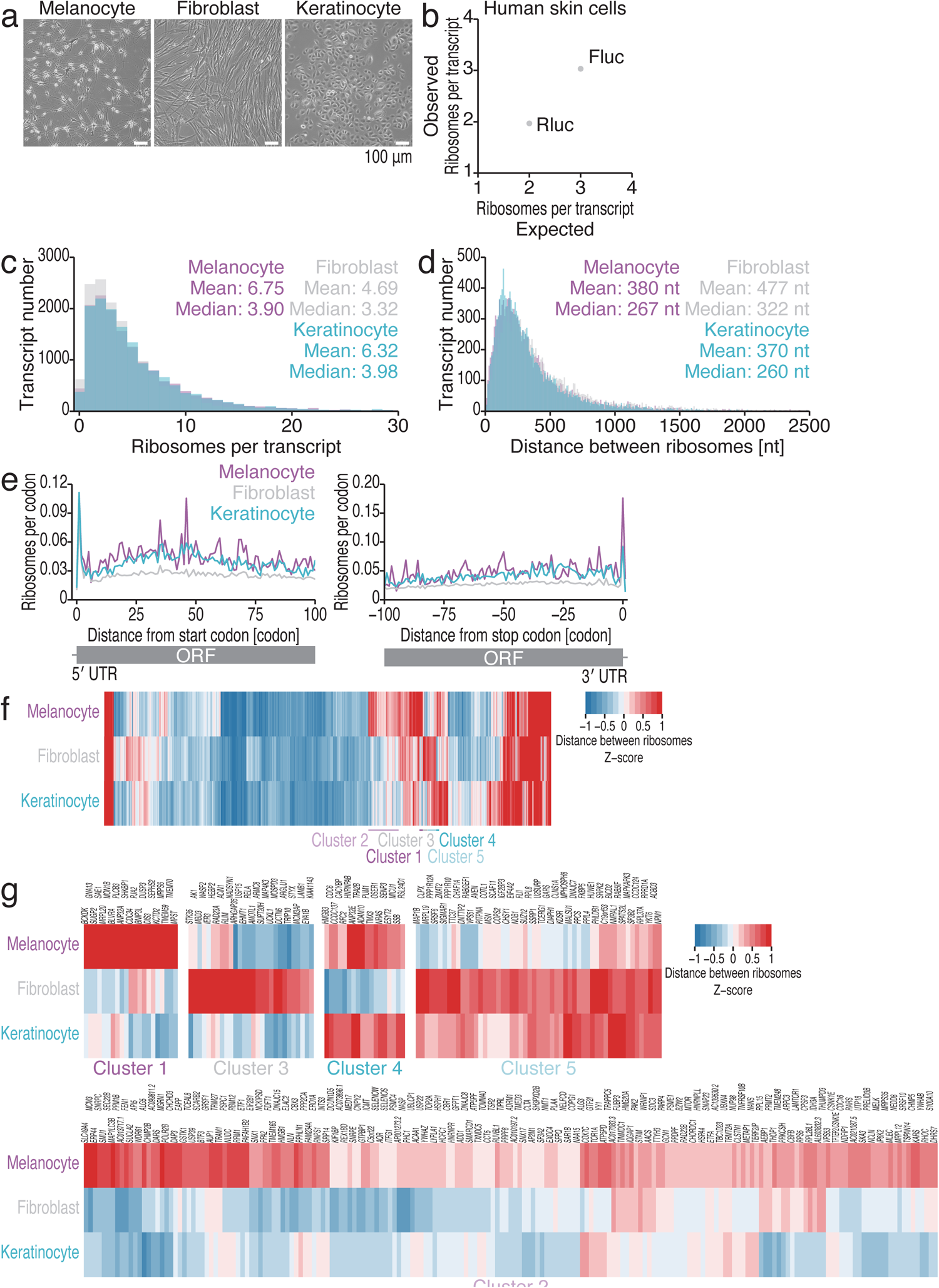
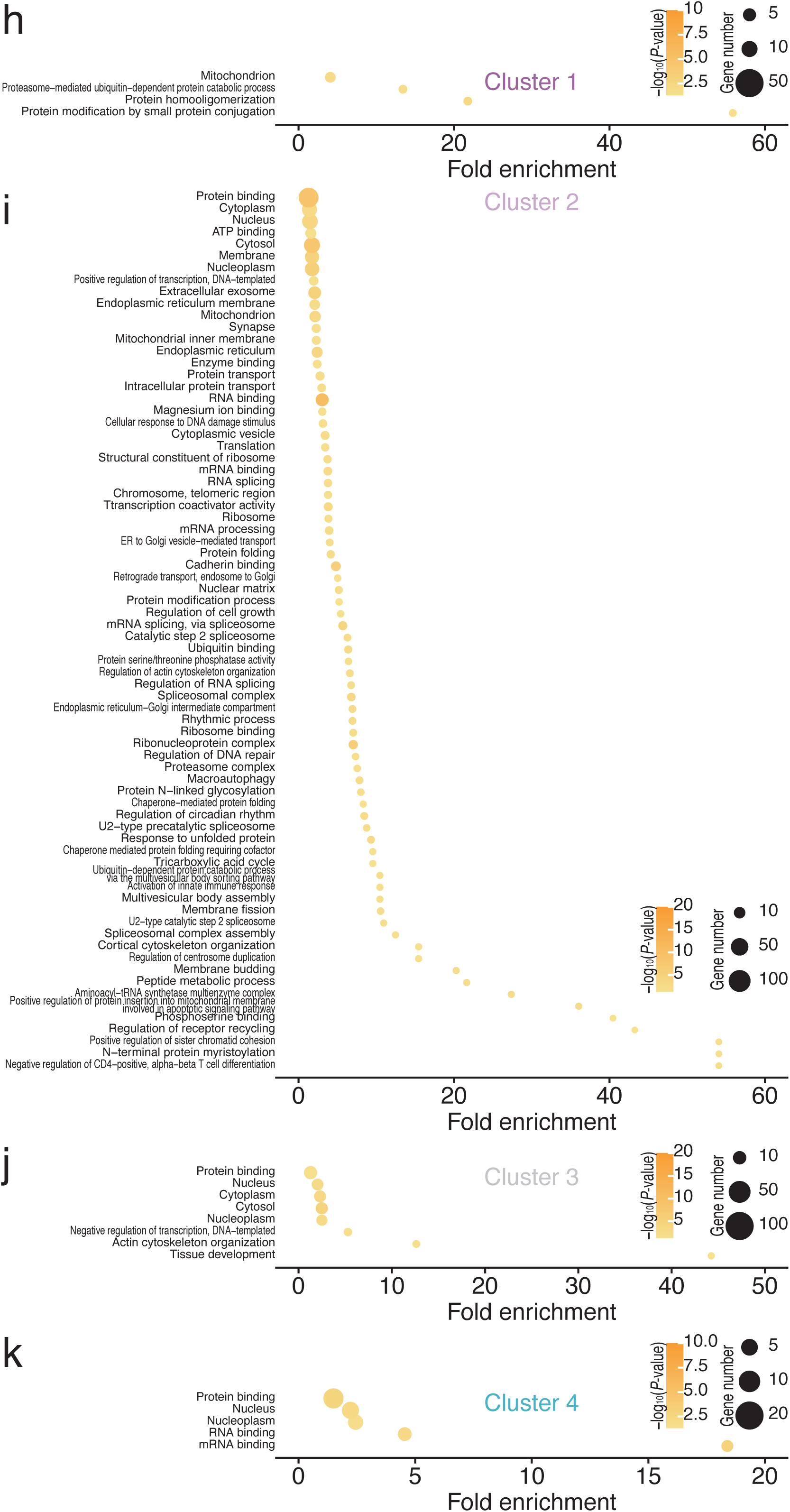
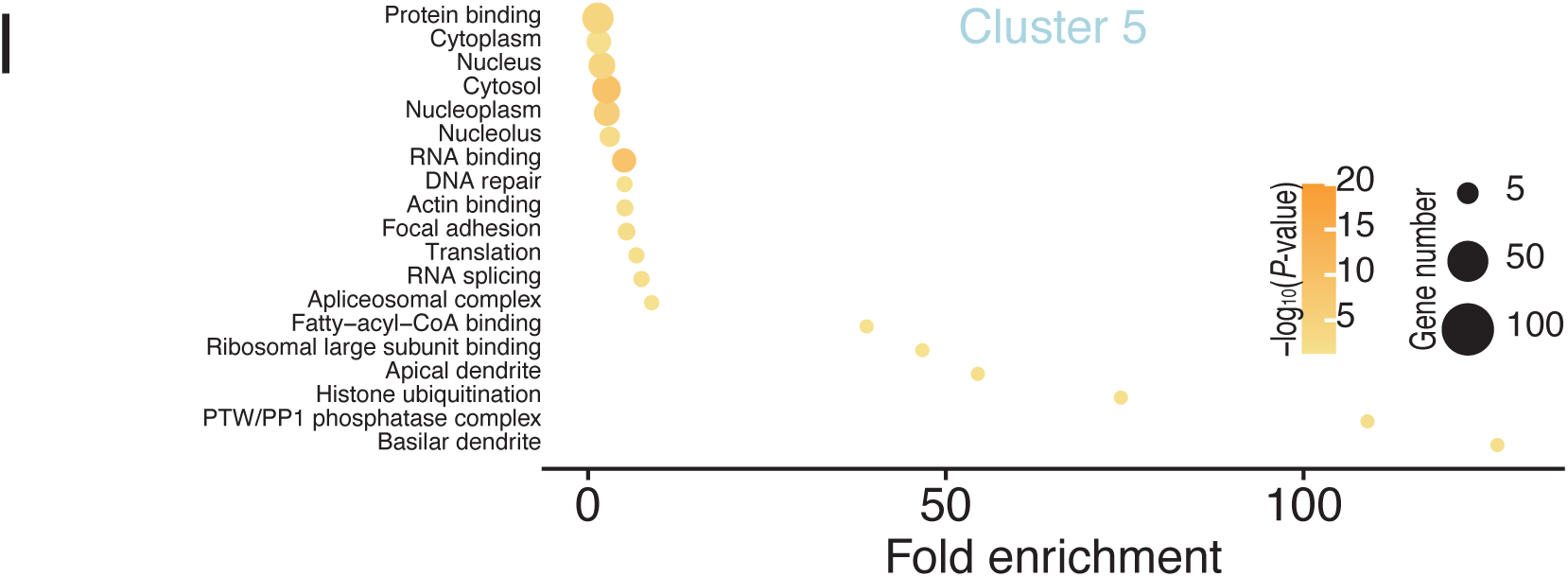
Characterization of ribosome profiling with Ribo-Calibration in skin cells. (a) The morphology of the indicated skin cells. Scale bar, 100 µm. (b) Comparison of expected and observed ribosome numbers on Rluc and Fluc transcripts used for Ribo-Calibration. (c) Histograms of the absolute ribosome number on transcripts in the indicated skin cells. (d) Histograms of the distance between ribosomes in the indicated skin cells. (e) Metagene plots with calibrated ribosome numbers at each position around start codons (left, the first position of the start codon was set to 0) and stop codons (right, the first position of the stop codon was set to 0) in the indicated skin cells. (f-g) Heatmap of the distance between ribosomes in the indicated skin cells. The color scales for the Z scores are shown. (h-l) GO terms enriched in the gene clusters defined in f. The color scales for significance and the size scales for the number of genes in each category are shown.

**Supplementary Fig. 9:**
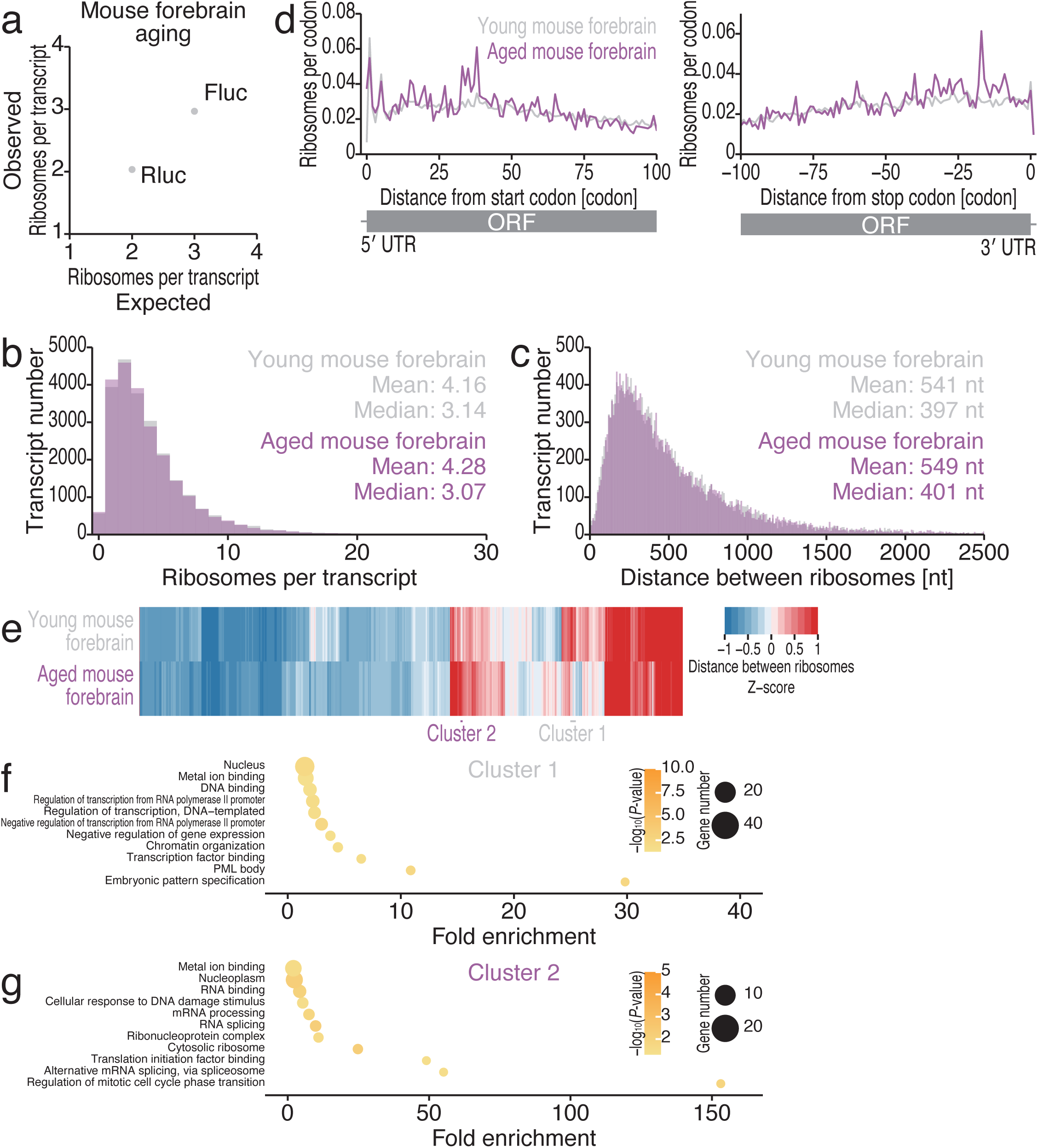
Characterization of ribosome profiling with Ribo-Calibration in mouse forebrains. (a) Comparison of expected and observed ribosome numbers on Rluc and Fluc transcripts used for Ribo-Calibration. (b) Histograms of the absolute ribosome number on transcripts in the young and aged mouse forebrains. (c) Histograms of the distance between ribosomes in the forebrains of young and aged mice. (d) Metagene plots with calibrated ribosome numbers at each position around start codons (left, the first position of the start codon was set to 0) and stop codons (right, the first position of the stop codon was set to 0) in the young and aged mouse forebrains. (e) Heatmap of the distance between ribosomes in young and aged mouse forebrains. The color scales for the Z scores are shown. (f-g) GO terms enriched in the gene clusters defined in e. The color scales for significance and the size scales for the number of genes in each category are shown.

